# Demographic history, geographic distance, and landscape features shape the genetic divergence of wild tigers in northeast India

**DOI:** 10.64898/2026.06.02.729668

**Authors:** Vinay Sagar, Tista Ghosh, Kritagnya Vadar, Vaishnavee Kundan Mayekar, Abishek Harihar, Uma Ramakrishnan

**Affiliations:** National Centre for Biological Sciences, TIFR, Bangalore 560065; Panthera, 104 W 40th St 5th Floor, NY, NY 10018; Nature Conservation Foundation, 1311,"Amritha", 12th Main, Vijayanagar 1st Stage, Mysore 570 017; Senior Fellow, Wellcome Trust DBT India Alliance

**Keywords:** genetic erosion, bottlenecks, carnivore connectivity, non-invasive genomics, fecal samples, conservation

## Abstract

Habitat fragmentation creates small, isolated populations vulnerable to inbreeding, genetic drift, and high genetic load. For conservation management, it is essential to distinguish contemporary landscape resistance from historical demographic processes as drivers of these genetic patterns, especially for conservation priority regions such as northeast India which intersects two major tiger conservation landscapes. We studied the genetic structure and landscape connectivity of tigers across four protected areas in northeast India: Kaziranga, Manas, Orang, and Nameri, using faecal samples. From 741 samples collected over two field seasons, we identified 654 confirmed tiger specimens. Using methylation-based enrichment and ddRAD-seq of 176 samples, we generated a high-quality dataset of 3091 SNPs across 44 individuals. Population structure analyses identified three genetically distinct clusters: Kaziranga-Nameri, Manas, and Orang. Isolation-by-distance and landscape resistance explained 13% and 17% of the observed genetic divergence, respectively, with human settlements influencing gene flow. Orang’s pronounced divergence from Kaziranga, despite geographic proximity and corridors, suggests a post-bottleneck founder effect, as evidenced by reduced heterozygosity (Ho = 0.24), nucleotide diversity (pi = 0.24), and effective population size (Ne = 1.3). These findings reveal that demographic and genetic recovery can decouple: Orang’s population has recently grown, yet genome-wide evidence shows ongoing genetic erosion that monitoring has not detected. Similar patterns have been reported in other Indian tiger populations, indicating that such decoupling may be systemic. Target 4 of the Kunming-Montreal Global Biodiversity Framework requires explicit genetic diversity monitoring; this study demonstrates that non-invasive genomics can operationalise that mandate at a conservation-relevant scale.

## 1. Introduction

Humans have altered ∼75% of the Earth’s ice-free land surface (Venter et al., 2016), with over 15% of the remaining natural habitats lost, and more than half of the world’s forests fragmented over the past few decades (Theobald et al., 2020; Zou et al., 2025). This trend is expected to intensify as ongoing habitat conversion and climate-driven land-use changes compound, accelerating the isolation of remnant populations across biomes (Zou et al., 2025). In addition to direct habitat loss, fragmentation disrupts population connectivity, making interpatch matrices hostile to dispersal, reducing effective population sizes, impeding gene flow, and elevating the risk of local extirpation (Crooks et al., 2017). Under such conditions, isolation amplifies genetic drift and inbreeding, causing erosion of adaptive genetic variation and stochastic fixation of harmful variants, often leading to demographic decline (Keller and Waller, 2002). Critically, such deterioration can persist even where structurally intact corridors seemingly link populations, as the translation of structural connectivity into realised genetic connectivity is mediated by behavioural, demographic, and landscape resistance factors (Correa Ayram et al., 2016; Lowe and Allendorf, 2010). Therefore, it is essential to disentangle landscape resistance from demographic history, both as a scientific priority and a policy requirement, directly linked to the Kunming-Montreal Global Biodiversity Framework’s Goal A, mandating species recovery to healthy and resilient levels, and Target 4, requiring explicit conservation of genetic diversity and adaptive capacity in wild populations (“CBD/COP/DEC/15/4,” 2022).

Meeting these objectives requires methods that can distinguish the contributions of contemporary landscape barriers from those of historical demographic processes. Landscape genetics addresses this need by integrating spatial statistics, landscape ecology, and population genetics to quantify the relative influence of environmental and anthropogenic features on gene flow (Manel et al., 2003). With advances in noninvasive genomics, such approaches are now feasible for wild, non-model species at scales relevant to conservation (Chiou and Bergey, 2018; Tyagi et al., 2022). However, a limitation of landscape genomic approaches is the risk of attributing genetic divergence primarily to current resistance or environmental gradients while underestimating the independent or confounding effects of demographic history (Dauphin et al., 2023). Isolated populations with small effective sizes remain susceptible to genetic drift, regardless of landscape configuration, as demonstrated in caribou (*Rangifer tarandus*), where drift-driven divergence persisted across seemingly connected habitats (Weckworth et al., 2013). Signals of past bottlenecks can persistently shape differentiation patterns for multiple generations because genetic responses lag landscape changes (Landguth et al., 2010). Hence, effective conservation planning requires a parallel evaluation of whether genetic structure reflects current resistance, historical demography, or their interaction, a distinction that has direct implications for population management and intervention urgency.

Tigers (*Panthera tigris*) are globally ’Endangered’ and ’Critically Depleted’, having lost over 90% of their historic range, with habitat fragmentation and loss of connectivity identified as primary impediments to functional recovery (Goodrich et al., 2022; Hunter et al., 2025). Low population densities, large home ranges, and sensitivity to human disturbance render the species disproportionately vulnerable to isolation (Ripple et al., 2014). Whole-genome analyses confirm that genetic drift is already the dominant force shaping variation in fragmented populations, with individual tigers carrying extended runs of homozygosity consistent with recent bottlenecks and founder events (Armstrong et al., 2021). India harbours ∼75% of the world’s tiger population and nearly half of its genetic variation (Mondol et al., 2009), but most populations persist in fragmented reserves with limited connectivity (Qureshi et al., 2023). The consequences are measurable; in a small, isolated reserve in western India (Ranthambore), the population carries elevated loads of deleterious variants consistent with inbreeding (Khan et al., 2021), whereas an isolated eastern Indian population (Simlipal) has undergone sufficient genetic drift to cause locally high frequency of the rare pseudomelanistic phenotype (Sagar et al., 2021). While corridors are expected to reduce extinction risk (Thatte et al., 2018), the assumption that structural corridors facilitate functional gene flow remains untested across most tiger landscapes.

Tigers in India’s northeastern region harbour unique genetic ancestry that may reflect historical gene flow with Indochinese populations, differentiating them from both Central and South Indian tigers (Kolipakam et al., 2019; Armstrong et al., 2021) and establishing them as a potential genetic bridge between the Eastern Terai and Indochinese conservation units (Jhala et al., 2021). The region, covering ∼295,000 km², encompasses two Tiger Conservation Landscapes (TCLs): Kaziranga TCL and the Indian portion of the Chittagong–Northern Triangle–Namdapha–Medog–Manas TCL. This area is one of the largest continuous tiger habitats, shared with Bangladesh, Bhutan, China, and Myanmar (Sanderson et al., 2023). Ensuring connectivity across this landscape is vital for demographic stability and sustained genetic exchange between countries, especially since the region is projected to support viable, functional tiger populations under optimistic recovery scenarios (Hunter et al., 2025). However, the Indian section is experiencing rapid and ongoing anthropogenic changes, especially in the Brahmaputra Valley, where built-up areas have increased by nearly 385% from 1973 to 2021 (Debnath et al., 2023). Connectivity is further threatened by hydroelectric development and land-use intensification (Pandit and Grumbine, 2012; Srinivasan et al., 2021). Previous genomic research has identified population sub-clustering among NE tigers despite seemingly intact structural corridors (Kolipakam et al., 2019), indicating a disconnect between structural and functional connectivity, the mechanistic basis of which remains unresolved.

In this study, we collected noninvasive faecal samples from four protected areas in this region and generated thousands of genome-wide SNP genotypes using methylation-based host DNA enrichment and ddRAD-seq. The four populations represent a structured demographic and connectivity gradient: Kaziranga Tiger Reserve, the main source population with the highest density and the most extensive protection history in the region; Nameri National Park, a small, low-density population connected to Kaziranga via Brahmaputra riverine corridors; Manas Tiger Reserve, a medium-sized, actively recovering population with functional transboundary connectivity to Bhutan’s Royal Manas (Lahkar et al., 2024); and Orang National Park, a small, high-density population surrounded by a highly modified matrix of agricultural and human settlements. We examined the following three questions: (1) the level of genetic differentiation among these populations and whether it correlates with geographic distance; (2) which landscape features most hinder gene flow and to what extent resistance explains divergence; and (3) where resistance is lacking, whether demographic history can account for residual structure. Owing to the demographic and connectivity gradient, and prior evidence of sub-clustering in regional tiger populations (Kolipakam et al., 2019), we hypothesised that the observed genetic structure would reflect both the imprint of contemporary landscape resistance and demographic history, with smaller populations embedded in rapidly changing landscapes expected to be most susceptible to drift-driven divergence. Our results are directly relevant to corridor conservation efforts in a landscape that significantly contributes to global tiger genetic diversity (Kolipakam et al., 2019; Armstrong et al., 2021).

## 2. Methods

### 2.1 Study Area

Tiger populations in northeastern India are distributed across four geographic blocks: (a) the Brahmaputra floodplain complex of Kaziranga, Karbi Anglong, Orang, Pakke, and Nameri, supporting over 150 tigers with densities exceeding 10 per 100 km² in Kaziranga and Orang; (b) the Manas-Buxa-Neora-Mahananda complex, supporting over 50 tigers, contiguous with Bhutan’s population of 131 tigers (Tempa et al., 2023); (c) the Dibang-Kamlang-Namdapha block in the northeast hills, characterised by low-density, scattered populations (Qureshi et al., 2023); and (d) the Dampa-Lushai hills region, where tiger presence is sporadic. Although forest corridors linking these blocks have been structurally identified (Qureshi et al., 2014), their functional connectivity is hindered by low population densities, intensive land-use change, and cultural variation among indigenous communities within intervening forest patches (Aiyadurai, 2016; Pandit and Grumbine, 2012; Srinivasan et al., 2021). Our sampling focused on four protected areas within blocks (a) and (b): Kaziranga Tiger Reserve, Nameri National Park, Manas Tiger Reserve, and Orang National Park.

### 2.2 Fieldwork and Sample Collection

Fieldwork and sample collection details can be found in the supplementary text S1.1.

### 2.3 Laboratory Processing and Genotyping

Details of DNA extraction, genetic species identification can be found in supplementary text S1.2 and S1.3. Faecal DNA extracts typically contain substantial bacterial DNA. Therefore, to enrich host DNA, we used the NEBNext Microbiome DNA Enrichment Kit (Cat. No. E2612L), following the protocols of and (Tyagi et al., 2022). ddRAD-seq Libraries were prepared as detailed in supplementary text S1.4. For cost-effectiveness, we downsampled before the enrichment and library preparation steps, as described in supplementary text S1.5.

### 2.4 Sequence data analysis pipeline, genetic differentiation and population structure

Details of sequence data analysis, variant discovery, variant filtering, unique individual identification and final dataset preparation are described in supplementary text S1.6 and S1.7, and population structure in S1.8.

### 2.5 Isolation by Distance and Landscape Resistance

Isolation by distance was tested in R (see supplementary text S1.9). *EEMS* (Petkova et al., 2016) was implemented to identify geographic barriers to gene flow beyond the IB, using merged district boundaries that encompassed all four protected areas as the outer boundary. Two independent MCMC chains were run with 200 demes, 5 million iterations, 50,000 burn-in, and 500 thinning steps; convergence was assessed from log-likelihood traces using *rEEMSplot*.

Landscape resistance was modelled using *ResistanceGA* (Peterman, 2018). Six variables were included: three natural—topographic position index (mTPI), NDVI, and permanent water body cover—and three anthropogenic—nightlight, built-up area proportion from the LULC layer, and linear infrastructure density from OpenStreetMap 2025 (details in supplementary text S1.9). Pairwise autocorrelation among variables was tested at r = 0.70. Variables were evaluated at six spatial resolutions (1, 2, 5, 10, 20, and 40 km) to assess scale dependence (Jackson and Fahrig, 2015). Each variable was optimised individually against pairwise genetic distance based on the proportion of shared alleles (DPS) using the MLPE approach and *SS_optim()* in *ResistanceGA*, with model selection via bootstrapping (*resist.boot()*, 1,000 iterations). The best-fitting univariate surfaces were combined into an optimised composite resistance surface using *MS_optim()*.

### 2.6 Population Genomic Diversity and Demographic History

Observed and expected heterozygosity (H_o_ and H_e_, nucleotide diversity (π), fixation index (F_IS_), pairwise F_ST_, and private and fixed allele counts/frequencies were estimated using the *Populations* module in *Stacks* (Catchen et al., 2013). Sample-size-corrected allelic richness was estimated by rarefaction in *ADZE-1.0* (Szpiech et al., 2008), and the distribution of major allele frequencies was examined across populations to quantify the extent of allele fixation consistent with drift. The effective population size was estimated using linkage disequilibrium-based methods in *NeEstimator V2* (Do et al., 2014).

## 3. Results

### 3.1 Sampling, Genotyping, and Dataset Summary

We collected 741 non-invasive samples (Kaziranga: 440, Manas: 197, Orang: 68, and Nameri: 36), of which 654 were genetically confirmed to be tigers (Kaziranga: 410, Manas: 158, Orang: 68, and Nameri: 18). Downsampling approach eliminated 249 samples before methylation-based enrichment, and 76 samples before ddRAD-seq library preparation, resulting in a final set of 329 ddRAD libraries. Among these, 153 libraries had concentrations too low to pool to the target concentration of 2 nM, resulting in a final pool of 176 sample libraries. Overall, 30.28% of all confirmed tiger samples were sequenced. We attempted to retain more samples from smaller populations (Nameri [83.33%], Orang [52.94%], Kaziranga [20.24%], and Manas [18.35%]; see Supplementary text S1.5 and Supplementary Table 1).

Sequencing produced an average of 11.1 million reads per sample, with a mean per-base quality score of 35.68 and >90% of base calls at a quality value of ≥30 across all samples. The average mapping rate per sample was 11.6% (Supplementary Table 4). Raw VCF contained 385,081,559 variant sites across 176 samples. After filtering, removing missing data, and excluding recaptured individuals, the final dataset comprised 44 unique individuals (Figure 1a, see Supplementary Figure S2 for recapture details) genotyped at 3,091 variant sites (Supplementary Table 2).

**Figure 1:**
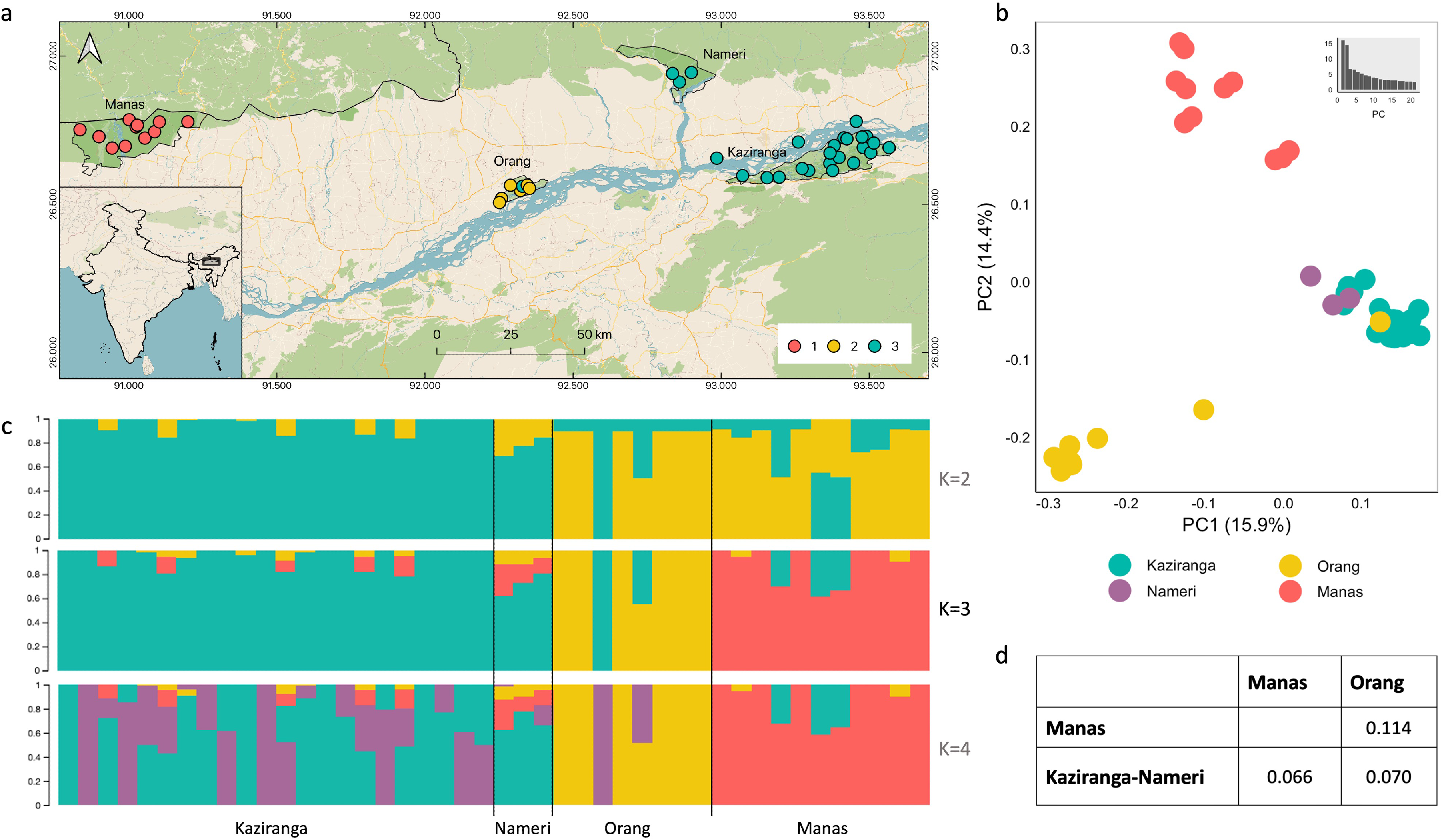
Tigers from Assam form three distinct genetic clusters: (1) Manas, (2) Kaziranga-Nameri, and (3) Orang. (a) Sampling location of 44 unique individuals coloured per their genetic clusters. The larger geographic distance of Manas from the other three PAs is noteworthy. (b) Principal Component Analysis identified three clusters. Nameri tigers group with Kaziranga tigers, while Manas and Orang tigers form their own separate groups. One individual from Orang is assigned to the Kaziranga cluster, suggesting dispersion from Kaziranga to Orang. (c) Bayesian clustering analysis supported the results from PCA; cross-validation error was minimum at K = 3, shown here is darker colour). One Orang individual derives ancestry from Kaziranga, supporting the PCA results. (d) Pairwise AMOVA F_ST_ for the three clusters; The strongest differentiation is observed between Manas and Orang.

### 3.2 Three genetically distinct tiger populations in the NE Brahmaputra floodplains

Principal component analysis (PCA) and Bayesian clustering (*Admixture*) identified three genetic clusters in the dataset. Manas and Orang tigers each formed distinct clusters, whereas Nameri and Kaziranga tigers were genetically indistinguishable (Figures 1b and 1c). *Admixture* analysis supported K=3 as the optimal number of clusters based on minimum cross-validation error. At K=4, Kaziranga tigers split into two groups broadly along an east-west axis, with Nameri continuing to group with Kaziranga (Supplementary Figure S1). Therefore, for all subsequent analyses, we treated three genetic populations: (1) Manas, (2) Kaziranga-Nameri, hereafter referred to as Kaziranga, and (3) Orang. Pairwise AMOVA F_ST_ values indicated that Kaziranga tigers were more similar to both Orang and Manas (Kaziranga-Orang F_ST_= 0.070; Kaziranga-Manas F_ST_ = 0.066; Figure 1d, Supplementary Figure S3) than Orang and Manas were to each other (Manas-Orang F_ST_ = 0.114).

### 3.3 Human settlements provide landscape resistance, yet explain only a minority of genetic divergence

The Mantel correlogram of isolation-by-distance analysis revealed a significant (p < 0.001) but weak correlation between genetic and geographic distance (adjusted R² = 0.128), with the relationship diminishing beyond approximately 50 km (Figure 2a). The kernel density plot revealed a discontinuous distribution with high-density clusters at specific distances, consistent with discrete barriers to gene flow in this landscape (Figure 2b). EEMS analysis supported this pattern, identifying major barriers to gene flow between Kaziranga and Orang, followed by Nameri and Manas (Figure 3c).

**Figure 2:**
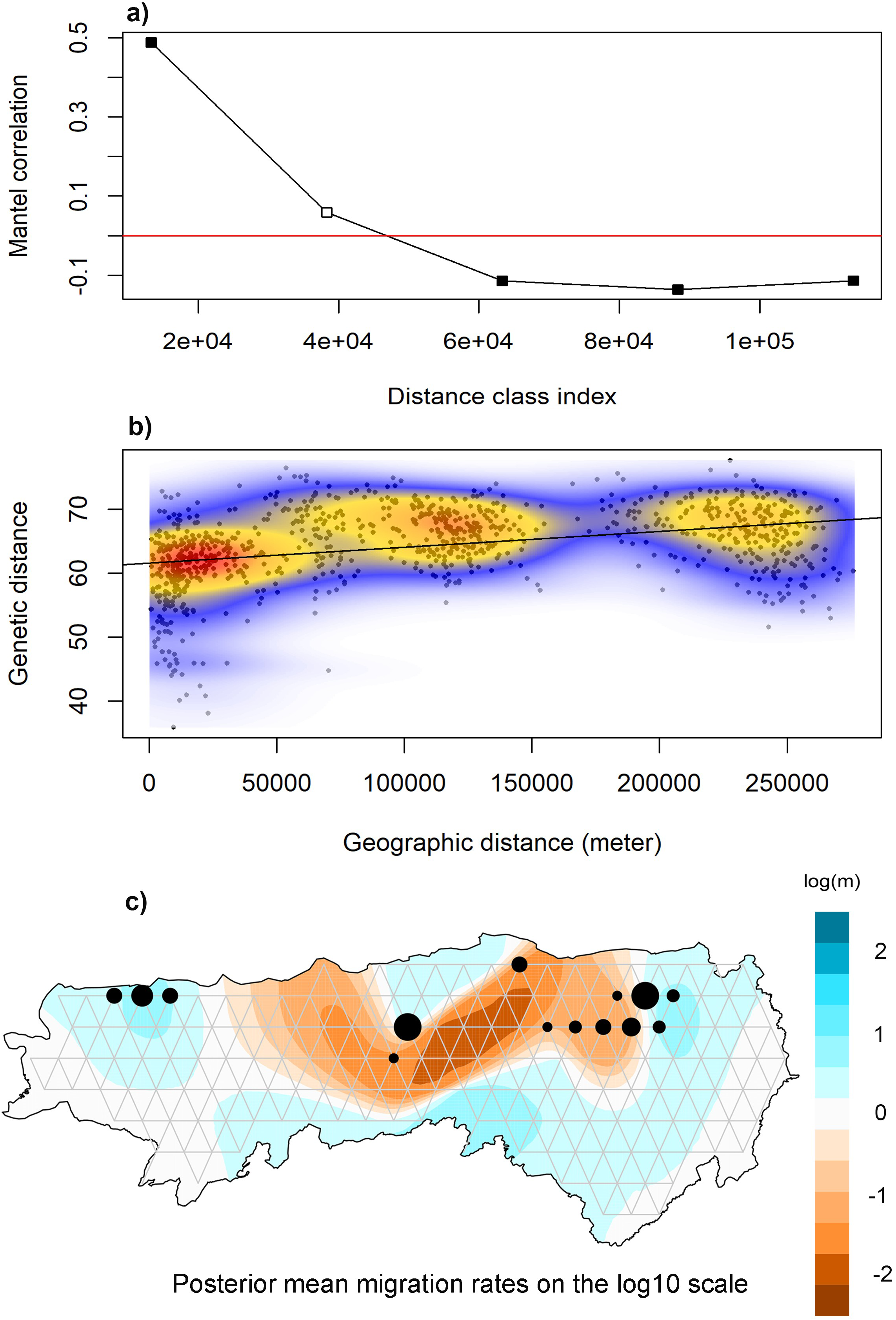
Testing isolation by distance and resistance. a) Mantel test of correlation between geographic and genetic distances. The solid points show the significant correlation between geographic and genetic distance where the red horizontal line is the threshold for no correlation (zero), (b) Geographic distance vs. genetic distance scatter plot with the linear relation shown with the black line. The point density decreases as the cloud colour changes from red to yellow to blue. (c) EEMS highlighting the areas impeding gene flow by estimating the disruption of migration rates in reaction to geographic distance, that is, isolation by resistance. The brown-blue colour gradient represents the rate of migration (low to higher than null expectation) between the sampling points, i.e., brown represents regions with physical barriers.

**Figure 3:**
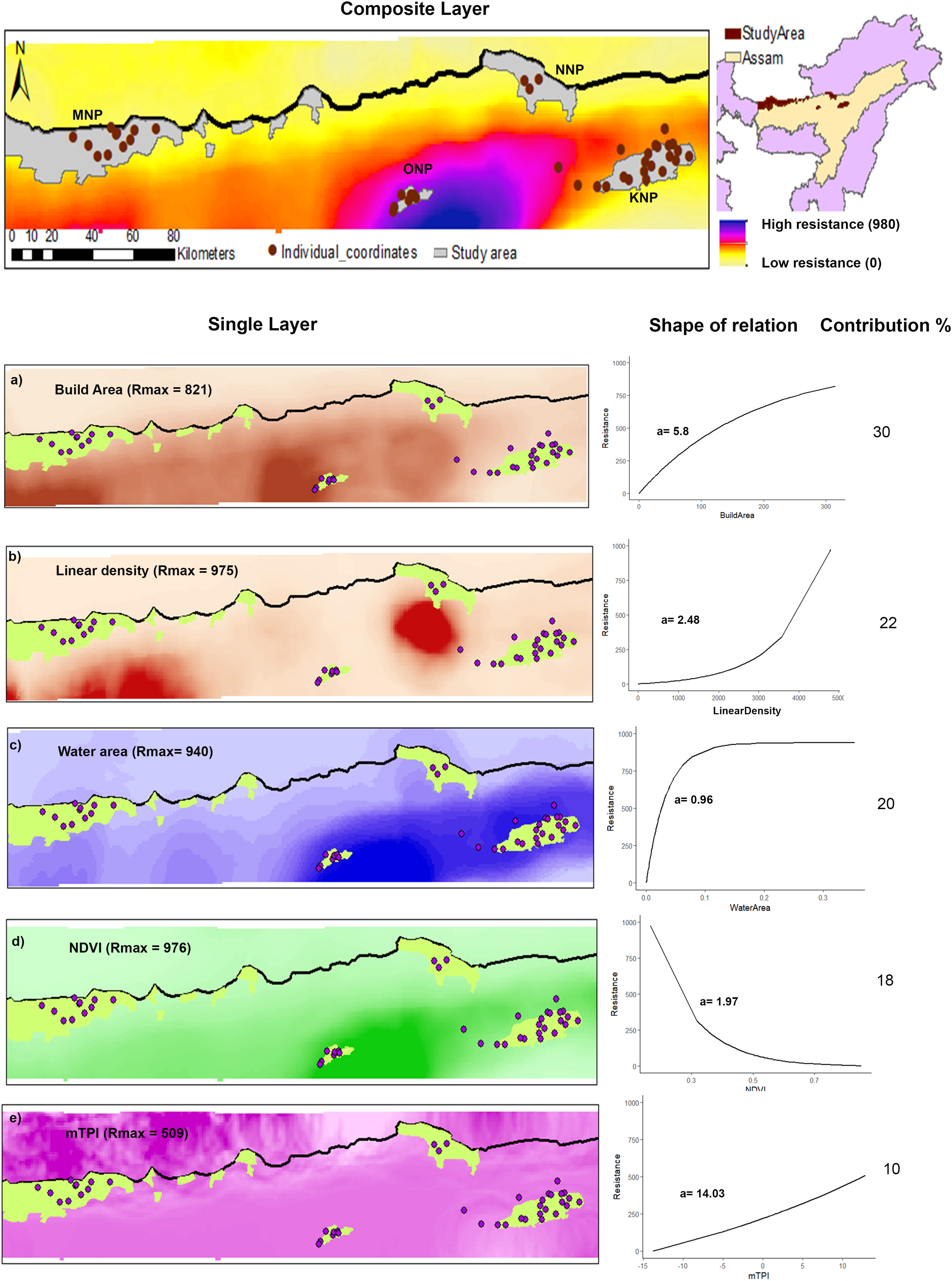
Identification of barriers to tiger gene flow across the study area. The upper panel shows the final composite resistance surface layer, where the dark-light red colour palette represents high-low resistance between the protected areas shown in green polygons. The lower panel (a-e) consists of selected individually optimised resistance surfaces arranged in decreasing order of their respective contributions (shown as contribution %) to the composite layer. For each independent layer, its Rmax and transformation graph were provided, showing the cost of animal movement across a particular landscape feature type. ‘a’ is the shape parameter that controls the rescaling of raw resistance values into movement costs required to optimise layers according to the observed genetic difference.

Landscape resistance modelling identified human settlements as the main factor hindering gene flow (Figure 3). Univariate optimization revealed spatial resolutions of 20 km for linear infrastructure density, built-up area, and permanent water bodies, and 10 km for NDVI and topography as the scales at which each variable most strongly predicted genetic distance (Supplementary Table 5). Among anthropogenic factors, built-up area had the most consistent effect, with resistance increasing asymptotically with settlement density (R_max_ = 821; Figure 3a). Linear infrastructure density showed higher maximum resistance than built-up area but only at high densities (R_max_ = 975). Among natural factors, NDVI had a high R_max_ of 976 but was limited to the narrow range of 0 to 0.3, representing barren and dry scrubland. Permanent water bodies (R_max_ = 940) were the most influential natural variable, offering consistently high resistance beyond 40 km². Topography contributed minimally, reflecting the broadly similar physiographic features of the study area. The final composite resistance surface explained 17% of the observed genetic variation (mR² = 0.174), with built-up area accounting for 30%, followed by linear infrastructure density, water body cover, NDVI, and topography at 22%, 20%, 18%, and 10%, respectively.

### 3.4 Convergent genomic signatures of isolation and drift in Orang tigers

Orang exhibited the lowest mean observed heterozygosity (H_O_ = 0.24) and mean nucleotide diversity (π = 0.24) among the three populations, compared with Kaziranga (H_O_ = 0.34; π = 0.32) and Manas (H_O_ = 0.33; π = 0.30; Table 1, Figure 4a and 4b). The distributions of H_O_ and π displayed a noticeable shift toward lower values in Orang. Notably, Manas had a higher proportion of SNPs with H_O_ and π values near zero than Kaziranga, despite similar means, suggesting a loss of low-frequency variation in Manas. The mean F_IS_ was negative in all three populations (Table 1), with Orang being the least negative (F_IS_ = -0.01) compared with Kaziranga (F_IS_ = -0.04) and Manas (F_IS_ = -0.06); however, the positive tail of the F_IS_ distribution gradually shifted toward higher values from Kaziranga through Manas to Orang, indicating increased inbreeding in Orang (Supplementary Figure S4).

**Figure 4:**
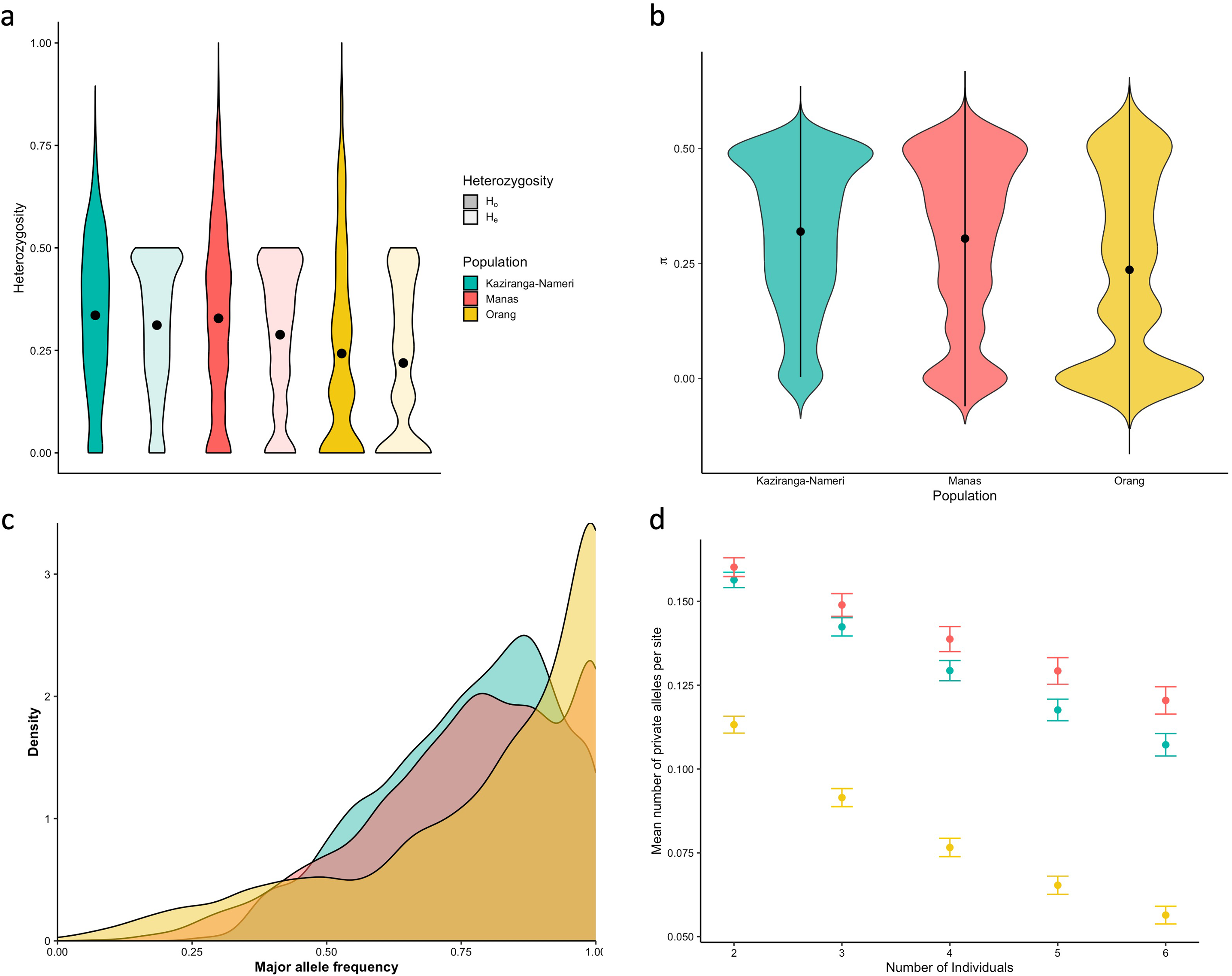
Reduced genetic variation in the Orang tigers. (a) Reduced heterozygosity in Orang compared to Manas and Kaziranga; the black dot shows the mean heterozygosity. Notably, all three clusters deviated from the Hardy-Weinberg Equilibrium, as evidenced by the differences between the expected and observed heterozygosity values (b) Violin plot showing the distribution of nucleotide diversity (π) in the three genetic clusters. While all populations showed reduced genetic diversity at several loci, Orang tigers showed the maximum loss. Low values of π also imply a low effective population size of Orang tigers. (c) A large fraction of alleles in all three populations showed a right shift from 0.5, indicating a drift towards the fixation of alleles. Orang showed the largest fraction of alleles that were fixed or reaching fixation, suggesting a strong effect of drift in the population. (d) Mean number of private alleles per site plotted against sample size after correcting for sample size bias. Orang had a significantly smaller number of private alleles than Manas and Kaziranga (p = 0.02).

**Table 1:**
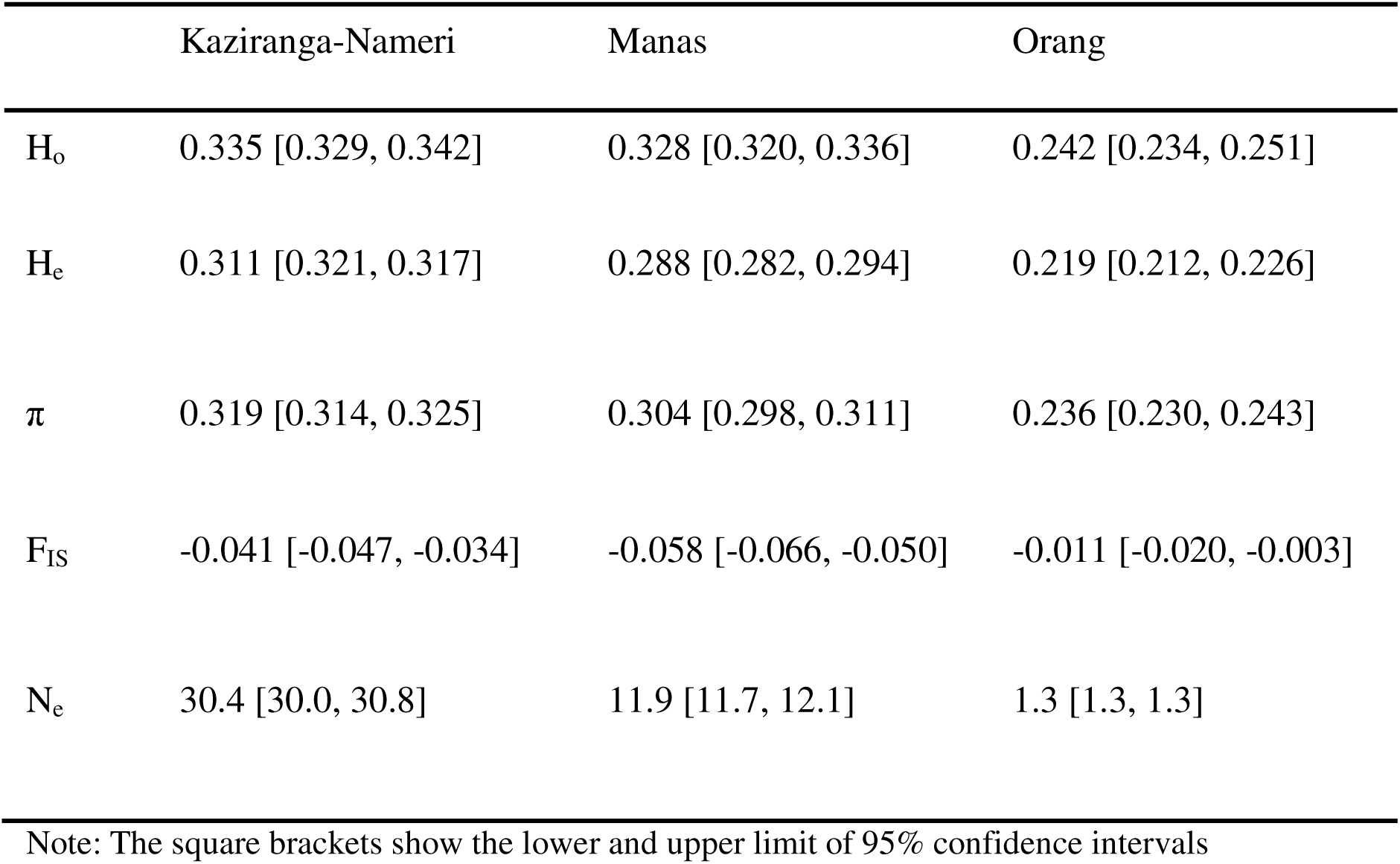
Population summary statistics for the three genetic clusters.

The distribution of major allele frequencies showed a significantly higher proportion of alleles fixed/nearly fixed in Orang compared to Kaziranga or Manas (Figure 4c). Rarefaction analysis indicated that the mean number of distinct alleles per site was lower in Orang than in either Kaziranga or Manas, although this difference was not statistically significant (Supplementary Figure S5; pairwise Wilcoxon test: Orang-Manas p = 0.33, Orang-Kaziranga p = 0.33, Manas-Kaziranga p = 0.69). The mean number of private alleles per site was significantly lower in Orang than in both Kaziranga and Manas (Figure 2D; pairwise Wilcoxon test: Orang-Manas p = 0.024, Orang-Kaziranga p = 0.024, Manas-Kaziranga p = 0.548), suggesting that alleles present in the broader NE tiger gene pool have been lost from Orang following population separation. The effective population size of Orang was estimated to be N_e_ = 1.3 (95% CI: 1.3–1.3), markedly lower than Kaziranga’s N_e_ = 30.4 (95% CI: 30.0–30.8) and Manas’s N_e_ = 11.9 (95% CI: 11.7–12.1; Table 1, Supplementary Table 3). Overall, Orang exhibited convergent signatures of a small, isolated population heavily shaped by genetic drift.

## 4. Discussion

The genetic structure of NE Brahmaputra tiger populations is shaped by both historical demographics and contemporary landscape resistance, and understanding their relative contributions is fundamental to assessing and managing these populations. Isolation by distance and landscape resistance explain ∼13% and 17% of genetic divergence, respectively, with human settlements identified as the primary barrier to gene flow. Notably, the pronounced divergence of Orang from Kaziranga, despite their geographic proximity and apparent structural connectivity, suggests a post-bottleneck founder effect and pervasive genetic drift. Consequently, landscape genetic models alone are insufficient to fully characterize the conservation status of this system.

### 4.1 Population Structure Does Not Reflect Geographic Proximity

As genetic differentiation and geographic distance do not correlate beyond a certain threshold, spatial proximity is not the primary driver of genetic connectivity in this landscape. The genetic indistinguishability of Nameri and Kaziranga, even at finer resolution (K = 4), is consistent with gene flow through their linking riverine corridor, reinforcing prior evidence of tiger movement along this route (Jhala et al., 2015). Notably, the Kaziranga-Orang F_ST_ (0.070) is similar to Kaziranga-Manas F_ST_ (0.066), despite the much greater distance to Manas, suggesting that the proximity between Kaziranga and Orang has not led to proportionally greater genetic exchange. The highest differentiation was observed between Manas and Orang (F_ST_ = 0.114), approaching the magnitude reported between South and Central Indian tiger populations separated by more than 1,500 km (Natesh et al., 2017), reinforcing that even modest F_ST_ values can reflect considerable effective isolation (Wright, 1949). The near absence of a direct structural corridor between Manas and Orang (Qureshi et al., 2014) provides a plausible structural explanation for this differentiation; however, as our results highlight, the absence of a structural corridor alone is insufficient to account for the full divergence pattern.

IBD analyses corroborate these observations. A significant but weak correlation between genetic and geographic distance (adjusted R² = 0.128) breaks down at approximately 50 km, well below the median predicted dispersal distance of 85 km (see Thatte et al., 2020). In the Central Indian landscape, where multiple populations show documented high gene flow (Schoen et al., 2022; Thatte et al., 2020), IBD breaks down at approximately 140 km, a directional contrast suggesting greater resistance to movement in northeast India. We caution that the Central India study used fewer microsatellite markers, known to influence IBD detection (Sgarlata et al., 2025). IBD explains only 12.8% of observed divergence indicating that much of the differentiation arises from non-linear processes beyond distance (de Flamingh et al., 2024). EEMS analyses support this pattern and reveal spatially heterogeneous migration rates and pronounced barriers to gene flow between Kaziranga and Orang. We note that F_ST_ estimates based on sample sizes, particularly for Orang and Nameri, are susceptible to upward bias, and all pairwise comparisons involving these populations should be interpreted with caution. Nevertheless, the convergent signal across PCA, Admixture, F_ST_, and IBD analyses strengthens confidence in the overall structural inference.

### 4.2 Human Settlements Dominate Landscape Resistance, but Contemporary Barriers Explain Only a Minority of Observed Divergence

The dominance of human settlements over infrastructure likely reflects the region’s socioeconomic structure, characterised by dense rural habitation and paddy-dominated agriculture. This pattern is reinforced by a 385% expansion of built-up areas in the Brahmaputra Valley between 1973 and 2021, alongside a concurrent loss of approximately 19.5% of vegetation cover (Debnath et al., 2023). It is also consistent with findings from other Indian tiger landscapes (Biswas et al., 2022; Thatte et al., 2018) and broader regional analyses showing that settlements exert a stronger influence on tiger distribution across northeast India than roads or railways (Deka et al., 2023). The effect of linear infrastructure increased sharply at higher densities, suggesting that continued infrastructure development could substantially impede movement for tigers and other wide-ranging mammals, as documented in the Central Indian landscape (Chakraborty et al., 2021; Thatte et al., 2020).

Beyond human influence, the Brahmaputra River exhibits high resistance to gene flow, which increases steadily with area and peaks at 40 km^2^. Although riverine grasslands and river islands were identified as structural corridors linking Kaziranga with both Orang and Nameri (Borah et al., 2010; Qureshi et al., 2014), our results indicate that only the Kaziranga–Nameri connection remains functional. Frequent and extensive flooding in Assam compounds this barrier effect, disrupting wildlife movement and necessitating refuge migration towards the Karbi Anglong hills, a route that is likely impeded by National Highway 715, which runs parallel to Kaziranga’s southern boundary and is associated with considerable tiger mortality from roadkill (Goswami et al., 2021; Qureshi et al., 2023).

The genetic evidence for movement of tigers through the Kaziranga-Nameri corridor despite villages along its length (Jhala et al., 2015), indicates that tigers can disperse through agricultural matrices when settlement density remains below a critical threshold. This aligns with broader evidence that dispersal in large carnivores is often less constrained by landscape conditions than habitat use within established home ranges (Mateo-Sánchez et al., 2015). The absence of comparable connectivity between Kaziranga and Orang reflects the heavily modified, high-disturbance character of the western Brahmaputra River islands (Borah et al., 2010) and highlights a meaningful distinction between the partial permeability of rural agricultural matrices and the near-impermeability of dense settlement matrices to tiger dispersal (Warrier et al., 2020). However, the continued functionality of the Kaziranga–Nameri corridor cannot be assumed, especially because recently established barriers may not yet produce detectable signs given known lag times between landscape change and genetic response (Landguth et al., 2010).

### 4.3 A Post-Bottleneck Founder Effect Is the Most Plausible Driver of Orang’s Genetic Divergence

Despite earlier identification of a potential corridor linking Kaziranga and Orang through Laokhowa Wildlife Sanctuary (Qureshi et al., 2014), the observed F_ST_ (0.070) indicates insufficient gene flow, likely reflecting processes beyond contemporary landscape resistance – a hypothesis supported by convergent genomic signatures. Orang tigers exhibited the lowest observed heterozygosity, lowest nucleotide diversity, elevated inbreeding, and the highest proportion of variant sites fixed or nearing fixation, even after correcting for sample size (Figure 4; Figures S4 and S5). Orang also showed the lowest number of private alleles and the highest number of alleles present in the combined Kaziranga–Manas gene pool but absent in Orang (Figures S6 and S7), indicating systematic drift-associated allele loss and fixation – characteristics of small, isolated populations. Although, the effective population size was estimated to be Ne = 1.3 (95% CI: 1.3–1.3; Supplementary Table 3), linkage disequilibrium-based Ne estimation is known to produce downward bias and artificially narrow confidence intervals at small sample sizes, and this figure should be treated as evidence of a very small effective size versus a precise estimate. In Orang, stable tiger habitat covers approximately 70 km², while ∼200 km² comprises largely Brahmaputra River islands and channels unviable for breeding (Borthakur et al., 2011). This constraint renders long-term demographic self-sufficiency structurally improbable, irrespective of current local densities.

Population monitoring data provide plausible mechanisms driving these genomic signatures. Despite high mortality, the Orang population recovered from seven individuals in 2008 to 28 by 2020 (Borthakur et al., 2011). This severe bottleneck followed by founder-led demographic expansion typically generates the genetic signatures observed here (England et al., 2003). While genomic studies have identified rangewide bottleneck signatures in Indian tigers (Armstrong et al., 2021), likely driven by colonial-era hunting (Rangarajan, 2003), this localized demographic history strongly implies a recent founder effect drove the marked genetic differentiation between the Orang and Kaziranga populations.

Two individuals sampled in Orang, derived 100% and 45% ancestry from the Kaziranga cluster (Figure 1c), and showed a PI-HAT (a relatedness measure) value of 0.5 with each other, consistent with a parent-offspring or full-sibling relationship, provided partial evidence of recent dispersal from Kaziranga and possible reproduction in Orang. However, such occasional immigration is insufficient to counteract drift in a small, recovering population. Although one migrant per generation has been suggested as a minimum threshold for reducing the harmful effects of drift, this relationship assumes an equilibrium between gene flow and drift, which is rarely applicable to recently diverged natural populations (Lowe and Allendorf, 2010). F_ST_ measured between recently diverged populations underestimates the realised loss of gene flow because drift has had insufficient time to reach equilibrium, and maintaining comparable allele frequencies requires substantially more than occasional immigrants, with estimates suggesting Nm >10 (Lowe and Allendorf, 2010). Lag-time considerations further complicate interpretation: simulation studies show that genetic responses to barrier establishment may require 1–15 generations to become detectable (Landguth et al., 2010). Additionally, as a population contracts, it may undergo a connectivity ‘Allee’ effect, in which movement rates decline precipitously at very low population and occupancy levels, despite the nominal presence of a permeable matrix (Zeigler and Fagan, 2014). The strong genetic separation between Orang and Kaziranga should be interpreted as the combined outcome of historical bottlenecks, founder effects, structural habitat constraints, and emerging landscape resistance operating across different timescales.

### 4.4 Research Priorities and Information Needs

Various research priorities emerge to address information gaps before making responsible, evidence-based management decisions on corridor prioritisation, genetic augmentation, or translocation, given the cross-sectional design’s limited temporal and mechanistic resolution. For Orang, effective immigration rates from Kaziranga cannot be estimated from a single dispersal event and require individual-based monitoring across multiple generations; the occupancy and permeability of the Laokhowa and Burachapori stepping-stone habitats (Qureshi et al. 2014) remain unassessed and represent the most tractable immediate priority, particularly given the rapid expansion of the human footprint around Tezpur (Baruah and Kar 2021). Whole-genome sequencing is required to determine whether adaptive variation has been compromised. Kaziranga itself should not be considered immune: the increasing frequency of the golden tiger phenotype, a rare recessive trait, may signal isolation-mediated drift (Kubala et al. 2019; Sagar et al. 2021), consistent with elevated runs of homozygosity in NE Indian tigers relative to Central Indian and Terai populations (Armstrong et al. 2021), warranting a comprehensive whole-genome assessment. Given lag times between landscape change and genetic response, for Kaziranga–Nameri, repeat sampling at five- to ten-year intervals would detect emerging differentiation before it constrains management options; for Kaziranga–Manas, the low pairwise FST (0.066) is consistent with recent historical gene flow, but coalescent-based modelling is needed to determine whether connectivity is current or declining. Securing stepping-stone habitats, including Sonai-Rupai Wildlife Sanctuary (Qureshi et al. 2014), is a priority now that genetic justification has been established, particularly given Manas’s link to genetically diverse Bhutan populations (Dhendup et al. 2023). Floodplain–hill connectivity and transboundary characterisation via Manas remain unresolved (Dhendup et al. 2023; Lahkar et al. 2024), and the non-invasive ddRAD-seq framework demonstrated here provides a scalable means to address all these priorities.

### 4.5 Conclusion

We show that demographic and genetic recovery can decouple in small, fragmented populations: the rapid expansion of the Orang tiger population is a conservation success by conventional metrics, yet genome-wide evidence is consistent with genetic erosion, a trajectory that density-based monitoring cannot detect. Such decoupling appears systemic rather than exceptional, with tigers in Ranthambore (Khan et al. 2021) and Simlipal (Sagar et al. 2021) showing analogous deterioration alongside census-level stability. Integrating genomic monitoring into national and state-level management frameworks is warranted. At broader scales, NE Indian tigers contribute disproportionately to within-species genetic diversity as a bridge between the Eastern Terai and Indochinese conservation units (Kolipakam et al. 2019; Jhala et al. 2021), and the Manas–Bhutan connection positions this landscape as a node of international genetic exchange (Dhendup et al. 2023). Target 4 of the Kunming–Montreal Global Biodiversity Framework requires explicit conservation of genetic diversity and adaptive capacity (CBD/COP/DEC/15/4 2022), a mandate that cannot be verified through census monitoring alone. We demonstrate that it can be operationalised through non-invasive genomics at a conservation-relevant scale, revealing insights that conventional monitoring would miss.

## Acknowledgements

We thank the Assam State Forest Department (permit nos. WL/FG.31/Technical Committee/2019/Pt dated 20/06/2020 and 26/08/2022; Office order 14 dated 13/01/2021); Assam State Biodiversity Board (no. ABB/Permission/2012/225 dated 06/11/2019) for permits and fieldwork support. We are grateful to the forest department field staff for their assistance and protection during sampling, and to our field volunteers Tikily Tayeng, Imran Rehman Momin, Mrunmay Desai, Divya Shrinivasan, Suraj Poojary, Jaya Norah, Yasung, and Samiran Sarkar. We thank Abhinav Tyagi for guidance on methylation-based enrichment and ddRAD-seq, Divyashree Rana for guidance on landscape genetics. We are grateful to Varun Goswami, Divya Vasudev, and Prachi Thatte for valuable discussions throughout the project. Sequencing was conducted at the NCBS sequencing facility, and we thank the facility manager, Awdhesh Pandit, for his assistance. Genomic analyses were performed on the computing cluster supported under project no. 12-R&D-TFR-5.04-0900, Department of Atomic Energy, Government of India. This project was implemented by the National Centre for Biological Sciences, TIFR, with Panthera using funding from the Robertson Foundation; V.S., K.V., and V.M. were supported by Panthera for the duration of their association with the project. AH contributed to funding acquisition through his role at Panthera and declares no competing interests regarding the results of this study.

## References

15/4 Kunming-Montreal Global Biodiversity Framework, 2022.

Aiyadurai, A., 2016. ′Tigers are Our Brothers′: Understanding Human-Nature Relations in the Mishmi Hills, Northeast India. Conserv. Soc. 14, 305. 10.4103/0972-4923.197614

Armstrong, E.E., Khan, A., Taylor, R.W., Gouy, A., Greenbaum, G., Thiéry, A., Kang, J.T., Redondo, S.A., Prost, S., Barsh, G., Kaelin, C., Phalke, S., Chugani, A., Gilbert, M., Miquelle, D., Zachariah, A., Borthakur, U., Reddy, A., Louis, E., Ryder, O.A., Jhala, Y.V., Petrov, D., Excoffier, L., Hadly, E., Ramakrishnan, U., 2021. Recent Evolutionary History of Tigers Highlights Contrasting Roles of Genetic Drift and Selection. Mol. Biol. Evol. 38, 2366–2379. 10.1093/molbev/msab032

Biswas, S., Bhatt, S., Sarkar, D., Talukdar, G., Pandav, B., Mondol, S., 2022. Assessing tiger corridor functionality with landscape genetics and modelling across Terai-Arc landscape, India. Conserv. Genet. 23, 949–966. 10.1007/s10592-022-01460-8

Borah, J., Ahmed, M.F., Sharma, P.K., 2010. Brahmaputra River islands as potential corridors for dispersing tigers: A case study from Assam, IndiaJ. Int. J. Biodivers. Conserv. 2, 350–358.

Borthakur, U., Barman, R.D., Das, C., Basumatary, A., Talukdar, A., Ahmed, M.F., Talukdar, B.K., Bharali, R., 2011. Noninvasive genetic monitoring of tiger (Panthera tigris tigris) population of Orang National Park in the Brahmaputra floodplain, Assam, India. Eur. J. Wildl. Res. 57, 603–613. 10.1007/s10344-010-0471-0

Catchen, J., Hohenlohe, P.A., Bassham, S., Amores, A., Cresko, W.A., 2013. Stacks: an analysis tool set for population genomics. Mol. Ecol. 22, 3124–3140. 10.1111/mec.12354

Chakraborty, P., Borah, J., Bora, P.J., Dey, S., Sharma, T., Lalthanpuia, Rongphar, S., 2021. Camera trap based monitoring of a key wildlife corridor reveals opportunities and challenges for large mammal conservation in Assam, India. Trop. Ecol. 62, 186–196. 10.1007/s42965-020-00138-x

Chiou, K.L., Bergey, C.M., 2018. Methylation-based enrichment facilitates low-cost, noninvasive genomic scale sequencing of populations from feces. Sci. Rep. 8, 1975. 10.1038/s41598-018-20427-9

Correa Ayram, C.A., Mendoza, M.E., Etter, A., Salicrup, D.R.P., 2016. Habitat connectivity in biodiversity conservation: A review of recent studies and applications. Prog. Phys. Geogr. Earth Environ. 40, 7–37. 10.1177/0309133315598713

Crooks, K.R., Burdett, C.L., Theobald, D.M., King, S.R.B., Di Marco, M., Rondinini, C., Boitani, L., 2017. Quantification of habitat fragmentation reveals extinction risk in terrestrial mammals. Proc. Natl. Acad. Sci. 114, 7635–7640. 10.1073/pnas.1705769114

Dauphin, B., Rellstab, C., Wüest, R.O., Karger, D.N., Holderegger, R., Gugerli, F., Manel, S., 2023. Re-thinking the environment in landscape genomics. Trends Ecol. Evol. 38, 261–274. 10.1016/j.tree.2022.10.010

de Flamingh, A., Alexander, N., Perrin-Stowe, T.I.N., Donnelly, C., Guldemond, R.A.R., Schooley, R.L., van Aarde, R.J., Roca, A.L., 2024. Integrating habitat suitability modeling with gene flow improves delineation of landscape connections among African savanna elephants. Biodivers. Conserv. 33, 3231–3252. 10.1007/s10531-024-02910-0

Debnath, J., Sahariah, D., Lahon, D., Nath, N., Chand, K., Meraj, G., Farooq, M., Kumar, P., Kanga, S., Singh, S.K., 2023. Geospatial modeling to assess the past and future land use-land cover changes in the Brahmaputra Valley, NE India, for sustainable land resource management. Environ. Sci. Pollut. Res. 30, 106997–107020. 10.1007/s11356-022-24248-2

Deka, J.R., Ali, Sk.Z., Ahamad, M., Borah, P., Gopi, G.V., Badola, R., Sharma, R., Hussain, S.A., 2023. Can Bengal Tiger (Panthera tigris tigris) endure the future climate and land use change scenario in the East Himalayan Region? Perspective from a multiple model framework. Ecol. Evol. 13, e10340. 10.1002/ece3.10340

Do, C., Waples, R.S., Peel, D., Macbeth, G.M., Tillett, B.J., Ovenden, J.R., 2014. NeEstimator v2: re-implementation of software for the estimation of contemporary effective population size (Ne) from genetic data. Mol. Ecol. Resour. 14, 209–214. 10.1111/1755-0998.12157

England, P.R., Osler, G.H.R., Woodworth, L.M., Montgomery, M.E., Briscoe, D.A., Frankham, R., 2003. Effects of intense versus diffuse population bottlenecks on microsatellite genetic diversity and evolutionary potential. Conserv. Genet. 4, 595–604. 10.1023/A:1025639811865

Goodrich, J., Wibisono, H., Miquelle, D., Lynam, A.J., Sanderson, E., Chapman, S., Gray, T.N.E., Chanchani, P., Harihar, A., 2022. Panthera tigris: The IUCN Red List of Threatened Species 2022 (No. e. T15955A21486201), The IUCN Red List of Threatened Species. International Union for Conservation of Nature and Natural Resources. 10.2305/IUCN.UK.2022-1.RLTS.T15955A214862019.en

Goswami, V.R., Vasudev, D., Joshi, B., Hait, P., Sharma, P., 2021. Coupled effects of climatic forcing and the human footprint on wildlife movement and space use in a dynamic floodplain landscape. Sci. Total Environ. 758, 144000. 10.1016/j.scitotenv.2020.144000

Hunter, L., Harihar, A., Miquelle, D., Gray, T.N.E., Goodrich, J., Bennet, E.L., Rosen, T., Linkie, M., Carlton, E., Grace, M.K., 2025. Panthera tigris (Green Status assessment). The IUCN Red List of Threatened Species 2025 (No. e. T15955A1595520252).

International Union for the Conservation of Nature and Natural Resources.

Jackson, H.B., Fahrig, L., 2015. Are ecologists conducting research at the optimal scale? Glob. Ecol. Biogeogr. 24, 52–63. 10.1111/geb.12233

Jhala, Y.V., Gopal, R., Mathur, V., Ghosh, P., Negi, H.S., Narain, S., Yadav, S.P., Malik, A., Garawad, R., Qureshi, Q., 2021. Recovery of tigers in India: Critical introspection and potential lessons. People Nat. 3, 281–293. 10.1002/pan3.10177

Jhala, Y.V., Qureshi, Q., Gopal, R., 2015. Status of Tigers, copredators & prey in India, 2014. National Tiger Conservation Authority, New Delhi, and Wildlife Institute of India, Dehradun, New Delhi.

Keller, L.F., Waller, D.M., 2002. Inbreeding effects in wild populations. Trends Ecol. Evol. 17, 19–23.

Khan, A., Patel, K., Shukla, H., Viswanathan, A., van der Valk, T., Borthakur, U., Nigam, P., Zachariah, A., Jhala, Y.V., Kardos, M., Ramakrishnan, U., 2021. Genomic evidence for inbreeding depression and purging of deleterious genetic variation in Indian tigers. Proc. Natl. Acad. Sci. 118, e2023018118. 10.1073/pnas.2023018118

Kolipakam, V., Singh, S., Pant, B., 2019. Genetic structure of tigers ( Panthera tigris tigris ) in India and its implications for conservation. Glob. Ecol. Conserv. 20, e00710. 10.1016/j.gecco.2019.e00710

Lahkar, D., Ahmed, M.F., Begum, R.H., Das, S.K., Sarma, H.K., Swargowari, A., Jhala, Y.V., Samad, I., Harihar, A., 2024. Post-conflict recovery of tigers (*Panthera tigris*) in a transboundary landscape: The case of Manas National Park, India. Biol. Conserv. 300, 110837. 10.1016/j.biocon.2024.110837

Landguth, E.L., Cushman, S.A., Schwartz, M.K., McKELVEY, K.S., Murphy, M., Luikart, G., 2010. Quantifying the lag time to detect barriers in landscape genetics. Mol. Ecol. 19, 4179–4191. 10.1111/j.1365-294X.2010.04808.x

Lowe, W.H., Allendorf, F.W., 2010. What can genetics tell us about population connectivity? Mol. Ecol. 19, 3038–3051. 10.1111/j.1365-294X.2010.04688.x

Manel, S., Schwartz, M.K., Luikart, G., Taberlet, P., 2003. Landscape genetics: combining landscape ecology and population genetics. Trends Ecol. Evol. 18, 189–197. 10.1016/S0169-5347(03)00008-9

Mateo-Sánchez, M.C., Balkenhol, N., Cushman, S., Pérez, T., Domínguez, A., Saura, S., 2015. A comparative framework to infer landscape effects on population genetic structure: are habitat suitability models effective in explaining gene flow? Landsc. Ecol. 30, 1405–1420. 10.1007/s10980-015-0194-4

Mondol, S., Karanth, K.U., Ramakrishnan, U., 2009. Why the Indian Subcontinent Holds the Key to Global Tiger Recovery. PLOS Genet. 5, e1000585.

Natesh, M., Atla, G., Nigam, P., Jhala, Y.V., Zachariah, A., Borthakur, U., Ramakrishnan, U., 2017. Conservation priorities for endangered Indian tigers through a genomic lens. Sci. Rep. 7, 9614. 10.1038/s41598-017-09748-3

Pandit, M.K., Grumbine, R.E., 2012. Potential Effects of Ongoing and Proposed Hydropower Development on Terrestrial Biological Diversity in the Indian Himalaya. Conserv. Biol. 26, 1061–1071. 10.1111/j.1523-1739.2012.01918.x

Peterman, W.E., 2018. ResistanceGA: An R package for the optimization of resistance surfaces using genetic algorithms. Methods Ecol. Evol. 9, 1638–1647. 10.1111/2041-210X.12984

Petkova, D., Novembre, J., Stephens, M., 2016. Visualizing spatial population structure with estimated effective migration surfaces. Nat. Genet. 48, 94–100. 10.1038/ng.3464

Qureshi, Q., Jhala, Y.V., Yadav, S.P., Mallick, A., 2023. Status of tigers, co-predators and prey in India, 2022 (No. ISBN 81-85496-92-7). National Tiger Conservation Authority, Government of India, New Delhi, and Wildlife Institute of India, Dehradun, New Delhi.

Qureshi, Q., Saini, S., Basu, P., Gopal, R., Raza, R., Jhala, Y., 2014. Connecting Tiger Populations for Long-term Conservation (No. TR2014- 02). National Tiger Conservation Authority & Wildlife Institute of India, Dehradun.

Rangarajan, M., 2003. India’s Wildlife History: An Introduction. The University of Chicago Press, Delhi, India.

Ripple, W.J., Estes, J.A., Beschta, R.L., Wilmers, C.C., Ritchie, E.G., Hebblewhite, M., Berger, J., Elmhagen, B., Letnic, M., Nelson, M.P., Schmitz, O.J., Smith, D.W., Wallach, A.D., Wirsing, A.J., 2014. Status and Ecological Effects of the World’s Largest Carnivores. Science 343, 1241484. 10.1126/science.1241484

Sagar, V., Kaelin, C.B., Natesh, M., Reddy, P.A., Mohapatra, R.K., Chhattani, H., Thatte, P., Vaidyanathan, S., Biswas, S., Bhatt, S., Paul, S., Jhala, Y.V., Verma, M.M., Pandav, B., Mondol, S., Barsh, G.S., Swain, D., Ramakrishnan, U., 2021. High frequency of an otherwise rare phenotype in a small and isolated tiger population. Proc. Natl. Acad. Sci. 118. 10.1073/pnas.2025273118

Sanderson, E.W., Miquelle, D.G., Fisher, K., Harihar, A., Clark, C., Moy, J., Potapov, P., Robinson, N., Royte, L., Sampson, D., Sanderlin, J., Yackulic, C.B., Belecky, M., Breitenmoser, U., Breitenmoser-Würsten, C., Chanchani, P., Chapman, S., Deomurari, A., Duangchantrasiri, S., Facchini, E., Gray, T.N.E., Goodrich, J., Hunter, L., Linkie, M., Marthy, W., Rasphone, A., Roy, S., Sittibal, D., Tempa, T., Umponjan, M., Wood, K., 2023. Range-wide trends in tiger conservation landscapes, 2001 - 2020. Front. Conserv. Sci. 4. 10.3389/fcosc.2023.1191280

Schoen, J.M., Neelakantan, A., Cushman, S.A., Dutta, T., Habib, B., Jhala, Y.V., Mondal, I., Ramakrishnan, U., Reddy, P.A., Saini, S., Sharma, S., Thatte, P., Yumnam, B., DeFries, R., 2022. Synthesizing habitat connectivity analyses of a globally important human-dominated tiger-conservation landscape. Conserv. Biol. 36, e13909. 10.1111/cobi.13909

Sgarlata, G.M., Maié, T., de Zoeten, T., Salmona, J., Rasteiro, R., Chikhi, L., 2025. The effect of habitat loss and fragmentation on isolation by distance and divergence. Proc. Natl. Acad. Sci. 122, e2410951122. 10.1073/pnas.2410951122

Srinivasan, U., Velho, N., Lee, J.S.H., Chiarelli, D.D., Davis, K.F., Wilcove, D.S., 2021. Oil palm cultivation can be expanded while sparing biodiversity in India. Nat. Food 2, 442–447. 10.1038/s43016-021-00305-w

Szpiech, Z.A., Jakobsson, M., Rosenberg, N.A., 2008. ADZE: a rarefaction approach for counting alleles private to combinations of populations. Bioinformatics 24, 2498–2504. 10.1093/bioinformatics/btn478

Tempa, T., Wangmo, S., Choijey, T., Sunar, S., Rigzin, K., Duba, D., Wangchuk, U., 2023. DoFPS 2023. Status of Tigers in Bhutan: The National Tiger Survey Report 2021–2022. Bhutan Tiger Center, Department of Forests and Park Services, Ministry of Energy and Natural Resources, Royal Government of Bhutan, Thimphu, Bhutan, Thimphu, Bhutan.

Thatte, P., Chandramouli, A., Tyagi, A., Patel, K., Baro, P., Chhattani, H., Ramakrishnan, U., 2020. Human footprint differentially impacts genetic connectivity of four wide-ranging mammals in a fragmented landscape. Divers. Distrib. 26, 299–314. 10.1111/ddi.13022

Thatte, P., Joshi, A., Vaidyanathan, S., Landguth, E., Ramakrishnan, U., 2018. Maintaining tiger connectivity and minimizing extinction into the next century: Insights from landscape genetics and spatially-explicit simulations. Biol. Conserv. 218, 181–191. 10.1016/j.biocon.2017.12.022

Theobald, D.M., Kennedy, C., Chen, B., Oakleaf, J., Baruch-Mordo, S., Kiesecker, J., 2020. Earth transformed: detailed mapping of global human modification from 1990 to 2017. Earth Syst. Sci. Data 12, 1953–1972. 10.5194/essd-12-1953-2020

Tyagi, A., Khan, A., Thatte, P., Ramakrishnan, U., 2022. Genome-wide single nucleotide polymorphism (SNP) markers from fecal samples reveal anthropogenic impacts on connectivity: case of a small carnivore in the central Indian landscape. Anim. Conserv. 25, 648–659. 10.1111/acv.12770

Venter, O., Sanderson, E.W., Magrach, A., Allan, J.R., Beher, J., Jones, K.R., Possingham, H.P., Laurance, W.F., Wood, P., Fekete, B.M., Levy, M.A., Watson, J.E.M., 2016. Sixteen years of change in the global terrestrial human footprint and implications for biodiversity conservation. Nat. Commun. 7, 12558. 10.1038/ncomms12558

Warrier, R., Noon, B.R., Bailey, L., 2020. Agricultural lands offer seasonal habitats to tigers in a human-dominated and fragmented landscape in India. Ecosphere 11, e03080. 10.1002/ecs2.3080

Weckworth, B.V., Musiani, M., DeCesare, N.J., McDevitt, A.D., Hebblewhite, M., Mariani, S., 2013. Preferred habitat and effective population size drive landscape genetic patterns in an endangered species. Proc. R. Soc. B Biol. Sci. 280, 20131756. 10.1098/rspb.2013.1756

Wright, S., 1949. The Genetical Structure of Populations. Ann. Eugen. 15, 323–354. 10.1111/j.1469-1809.1949.tb02451.x

Zeigler, S.L., Fagan, W.F., 2014. Transient windows for connectivity in a changing world. Mov. Ecol. 2, 1. 10.1186/2051-3933-2-1

Zou, Y., Crowther, T.W., Smith, G.R., Ma, H., Mo, L., Bialic-Murphy, L., Potapov, P., Gawecka, K.A., Xu, C., Negret, P.J., Lauber, T., Wu, Z., Rebindaine, D., Zohner, C.M., 2025. Fragmentation increased in over half of global forests from 2000 to 2020. Science 389, 1151–1156. 10.1126/science.adr6450

