## Supplemental Information for "Demographic history, geographic distance, and landscape features shape the genetic divergence of wild tigers in northeast India"

#### Supplementary Information

### S1. Supplementary methods:

S1.1 Fieldwork and sample collection:

Sampling was conducted over two field seasons: January 15 to March 15, 2021, and December 22, 2022, to March 10, 2023. In all protected areas except Kaziranga, surveys were conducted on foot; in Kaziranga, where high-density Indian one-horned rhinoceros (*Rhinoceros unicornis*) populations posed a safety constraint, surveys were conducted by vehicle at less than 10 km/h speed. Sampling tracks were designed to maximise individual capture within each protected area, with mean track lengths of 15.4 km in walkable areas and 32.6 km in Kaziranga, based on tiger presence data from the Assam State Forest Department. Each track was surveyed twice per season with a minimum of six days between surveys. The total survey effort across both seasons was 547 and 540 km in Kaziranga, 239 and 259 km in Manas, 138 and 132 km in Orang, and 150 and 94 km in Nameri, yielding a combined effort of 4,198 km. Supplementary surveys of the Brahmaputra River islands across both seasons added 105 km of coverage, broadening spatial sampling. Faecal samples were collected by swabbing fresh scats with sterile polyester swabs stored in Longmire's buffer (Longmire et al., 1997); shed hair from scratch marks was collected in dry zip-lock bags. All samples were stored at room temperature until the end of each field season and then at -20°C. Sampling was conducted under permits from the Assam State Forest Department and the Assam State Biodiversity Board.

S1.2 DNA extraction:

DNA was extracted from faecal samples using Qiagen DNA extraction reagents as follows: 20 µL proteinase K (Cat. No. 19134) was added to 300 µL of the sample (dissolved in the sampling buffer) and incubated at 56°C for 10–12 h, followed by adding 400 µL of buffer AL (Cat. No. 19075) and incubating at 56°C for 10–20 min, and then adding 400 µL of 100% ethanol before loading on the Qiagen spin column. Two washing steps with buffer AW1 (Cat. No. 19081) and AW2 (Cat. No. 19072) were performed as recommended by the manufacturer. Finally, DNA was eluted in 200 µL nuclease-free water after 15–30 min at room temperature, followed by a final spin at 13,000 rpm for 1 min.

S1.3 Genetic species identification:

Species identification was performed by PCR amplification of a mitochondrial cytochrome B fragment using tiger-specific primers [TigCytB162bp; (Bhagavatula and Singh, 2006); 94°C/30 s, 62°C/30 s, 72°C/30 s, 40 cycles], confirmed by agarose gel electrophoresis. Samples that failed this assay were tested using a felid-specific primer to amplify a 202 bp mitochondrial 16S-rRNA fragment [16S-rRNA-Cat; (Sagar et al., 2021)] under identical cycling conditions, sequenced on the Sanger platform, and identified by NCBI-BLAST. Nuclease-free water and tiger tissue DNA served as negative and positive controls, respectively.

S1.4 ddRAD-seq library preparation:

The libraries were prepared following the approach of (Tyagi et al., 2022) as follows: simultaneous digestion with MluCI and SphI restriction enzymes (NEB, Cat. No. R0538L and R3182L), adaptor ligation, purification with AMPure XP beads (Product No: A63881), and dual-barcode attachment via four-replicate indexing PCR to minimise amplification bias. Dual-size selection targeting 200–300 bp fragments was performed using AMPure XP beads. Individual libraries were quantified using Qubit 2.0, size-verified using the High-sensitivity D1000 screentape on a Agilent 4200 TapeStation, pooled to an equimolar concentration of 2 nM, and sequenced on an Illumina NovaSeq 6000 (S2 flow cell).

S1.5 Downsampling approach:

To avoid enriching low-yield samples likely to fail this step, we first downsampled them using a 123-locus multiplex PCR panel (Natesh et al., 2019), sequenced them on an Illumina MiSeq (150-cycle V3 kit), and genotyped them as described by (Sagar et al., 2021). Samples with at least one technical replicate genotyped at more than 20 of the 123 loci were selected for enrichment. All samples from Orang and Nameri were included, regardless of genotyping performance, given their small sizes. The DNA concentration was quantified before and after enrichment using a Qubit 3.0 fluorometer (Cat. No. Q33216).

A pilot library of 28 samples (concentration range: below the detection limit to 0.8 ng/μL on the Qubit high-sensitivity assay) was prepared, and eight libraries were sequenced to calibrate mapping efficiency against the input concentration. Samples achieving at least 30% mapping efficiency had post-enrichment concentrations ≥ 0.01 ng/μL; this threshold was applied to all subsequent library preparations, with all Nameri samples included regardless of concentration, given the small sample sizes. Technical replicates were pooled prior to library preparation to maximise input DNA.

S1.6 Sequence data analysis pipeline and variant discovery:

Raw fastq reads were trimmed using fastp [(Chen, 2023); -n 5 -q 15 -u 30 --detect_adapter_for_pe --cut_right --cut_right_window_size=4 --cut_right_mean_quality=15 --length_required=25] and aligned to the tiger reference genome PanTigT.MC.v3 [(Shukla et al., 2023); GenBank: GCA_021130815.1] using bwa-mem2 (Li and Durbin, 2009). SAM files were coordinate-sorted, converted to BAM, and indexed using SAMtools (Danecek et al., 2021; Li et al., 2009); read groups were assigned with Picard AddOrReplaceReadGroups ("Picardtools," 2019). Variant calling was performed with BCFtools (Danecek et al., 2021), and the resulting VCF was filtered to remove low-quality calls (QUAL ≤10), low depth (INFO/DP ≤200), and sites with mapping quality bias (MQBZ <-3), read position bias (RPBZ <-3 or >3), strand bias (FORMAT/SP >30), soft-clip bias (SCBZ >3), or low mapping quality (MQ ≤30). Further filtering with VCFtools (Danecek et al., 2011) removed indels, retained only biallelic SNPs, and applied thresholds of minDP 10, minQ 30, minGQ 30, and MAF 0.05. Individuals with fewer than 2,000 genotyped sites and sites missing in more than 30% of individuals were removed, and a Hardy-Weinberg equilibrium filter (--hwe 0.05) was applied.

S1.7 Individual Identification and Dataset Preparation:

Duplicate samples were identified using the KING duplicate algorithm (Manichaikul et al., 2010), and samples with more missing data were removed from each duplicate pair. KING (--related --degree 2) was then used to identify highly related pairs; one sample showing anomalously high relatedness to multiple individuals, likely reflecting excessive missing data, was removed. Pairwise genetic relatedness (PI-HAT) was estimated using PLINK 1.9 (Chang et al., 2015; Purcell et al., 2007), and one sample from each pair with PI-HAT >0.7 was excluded to eliminate potential residual duplicates.

S1.8 Genetic Differentiation and Population Structure

Principal component analysis was performed using PLINK 1.9 (Chang et al., 2015). Bayesian clustering was conducted in ADMIXTURE (Alexander and Lange, 2011) for K = 2–5, with 10 independent runs per K; the optimal K was selected as the K with the minimum cross-validation error, and the results were summarised using CLUMPAK (Kopelman et al., 2015).

S1.9 Isolation by Distance Analysis:

Isolation by distance was tested in R using *adegenet* (Jombart, 2008; Jombart and Ahmed, 2011), with geographic distances calculated from UTM coordinates converted using the *sf* package (Pebesma, 2018). Mantel correlograms and kernel density plots of genetic versus geographic distance were generated using *vegan* (Oksanen et al., 2025) and *MASS* (Venables and Ripley, 2002), respectively.

S1.10 Landscape resistance layers:

Landscape resistance was modelled including six variables: three natural—topographic position index [mTPI; (Theobald et al., 2015)], NDVI (Didan, 2021), and permanent water body cover (Pekel et al., 2016)—and three anthropogenic—nightlight (Elvidge et al., 2021), built-up area proportion from the LULC layer (Brown et al., 2022), and linear infrastructure density from OpenStreetMap 2025. All layers except linear infrastructure were obtained from the Google Earth

Engine; built-up area was reclassified to a continuous proportion variable on a 1 km grid, and linear infrastructure was converted to linear density.

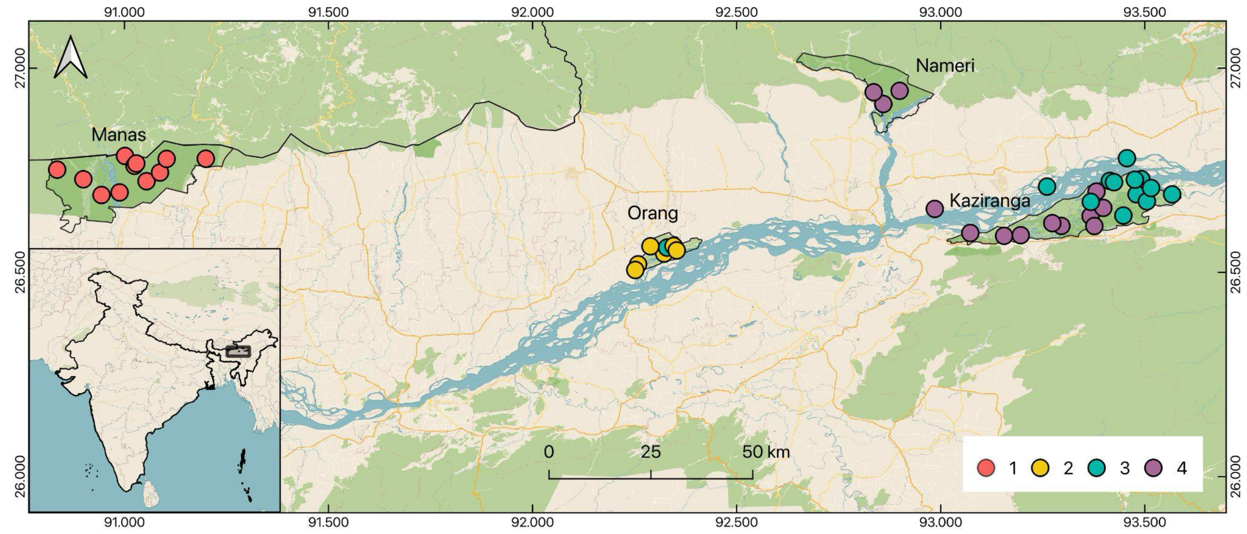


Figure S1: The genetic clusters obtained from *Admixture* at K=4 plotted on the landscape map. At K=4, Kaziranga tigers split into two genetic clusters, largely along the East - West axis. For this figure, the individuals were assumed to belong to a cluster if their shared ancestry in the Q-matrix was more than 0.5 in the given cluster.

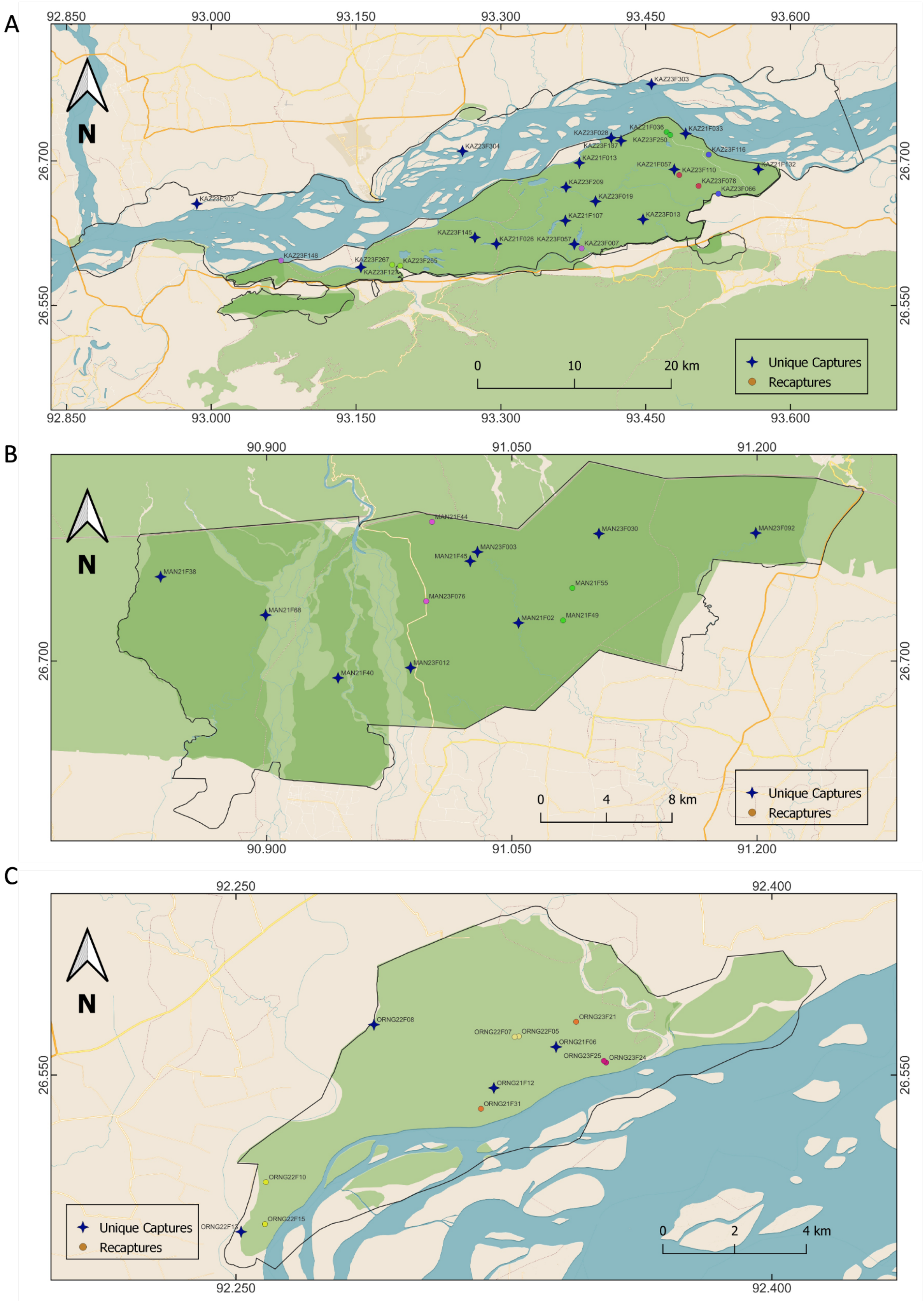


Figure S2: Recaptures in (A) Kaziranga, (B) Manas, and (C) Orang. The same individuals recaptured are shown in matching colours and non-recaptured individuals as blue stars.

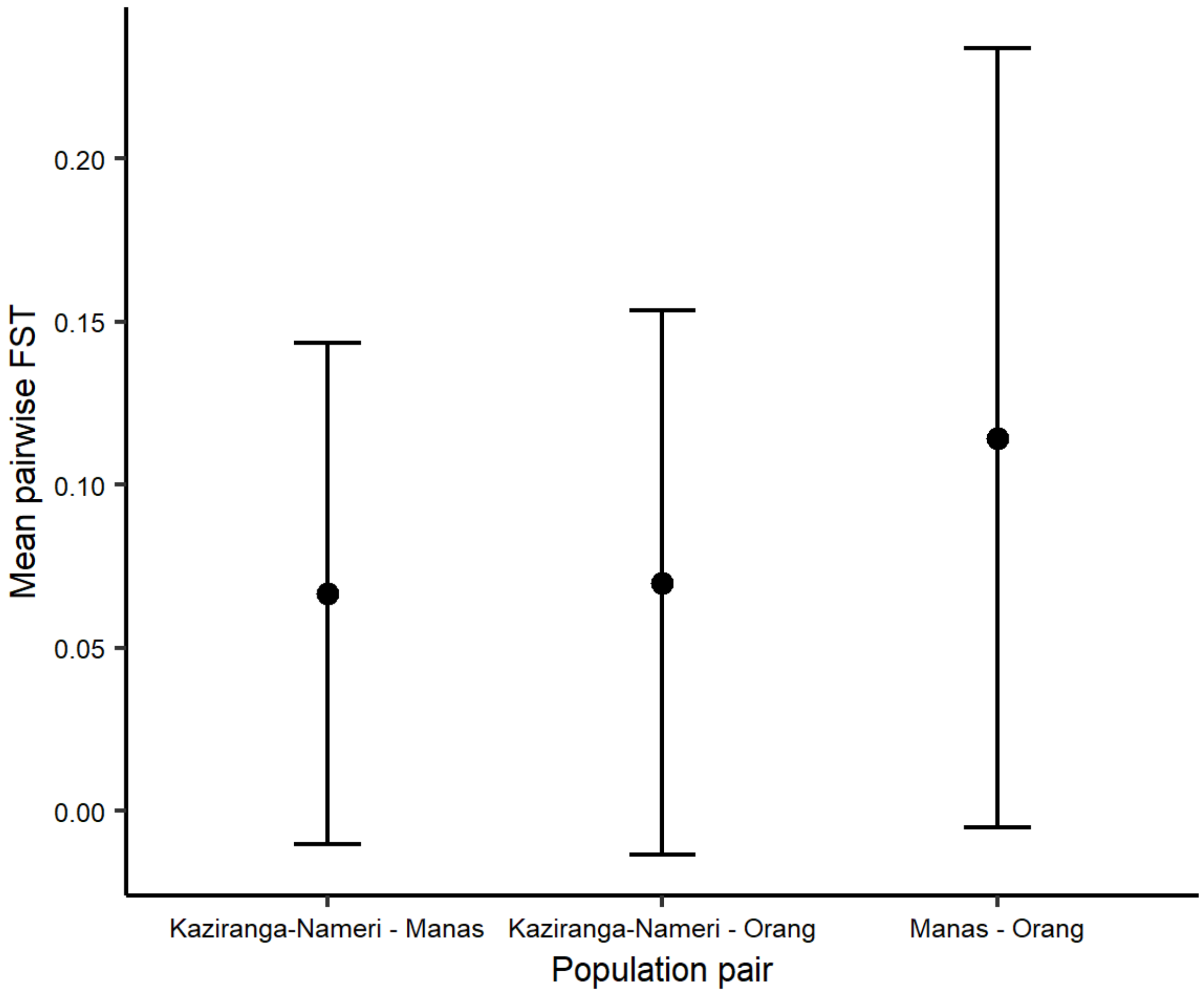


Figure S3: Mean pairwise AMOVA-$F_{ST}$ plot for all the sites. The error bars represent the standard deviation. Highest values were observed for Orang – Manas pair, while FST between Kaziranga – Orang and Kaziranga – Manas pairse was comparable.

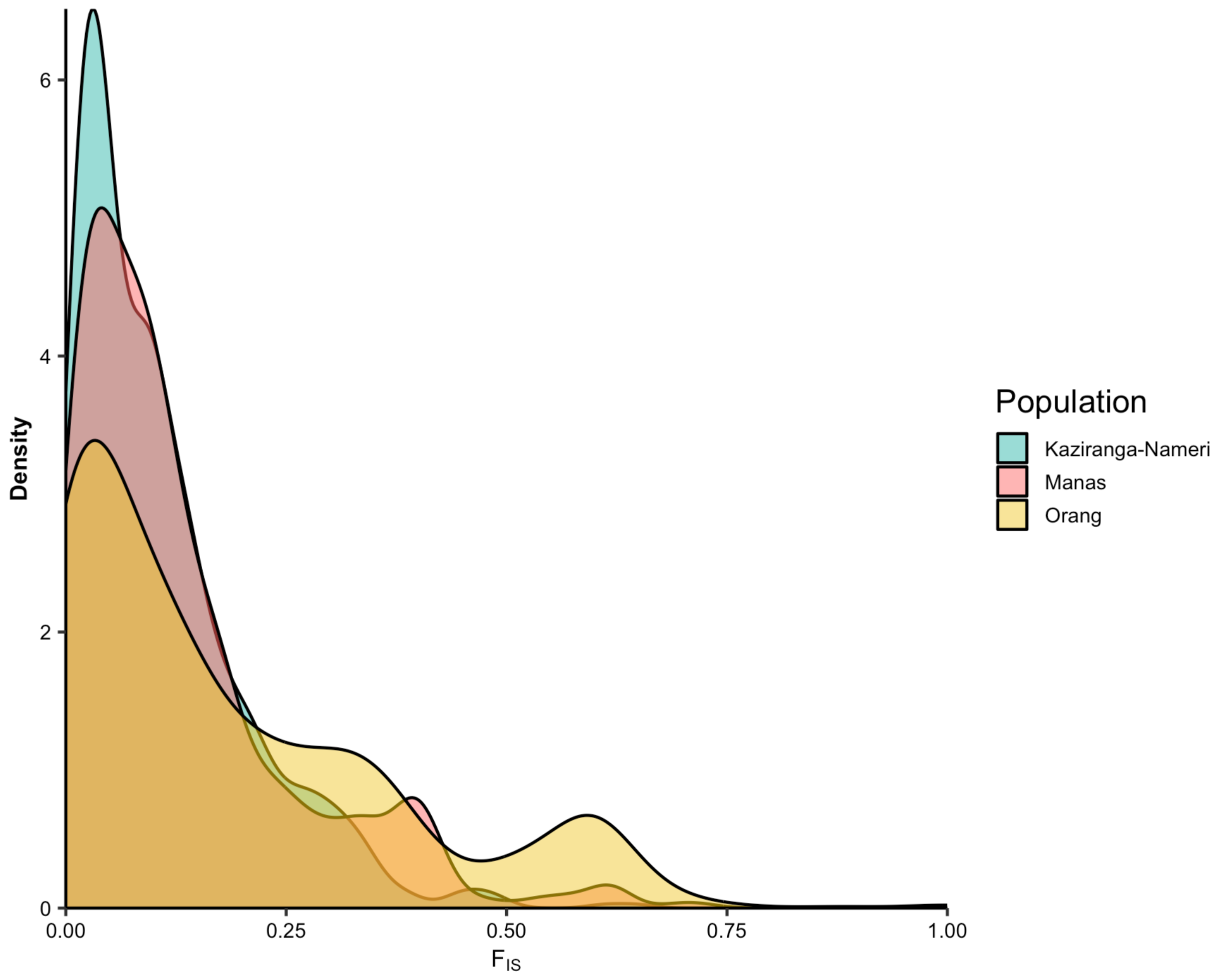


Figure S4: Distribution of positive fraction of $F_{IS}$ showing inbreeding in the three genetic clusters. Most loci in Kaziranga show low FIS values while a large fraction of loci in Orang show high FIS values, suggesting an increase in inbreeding in Orang tigers.

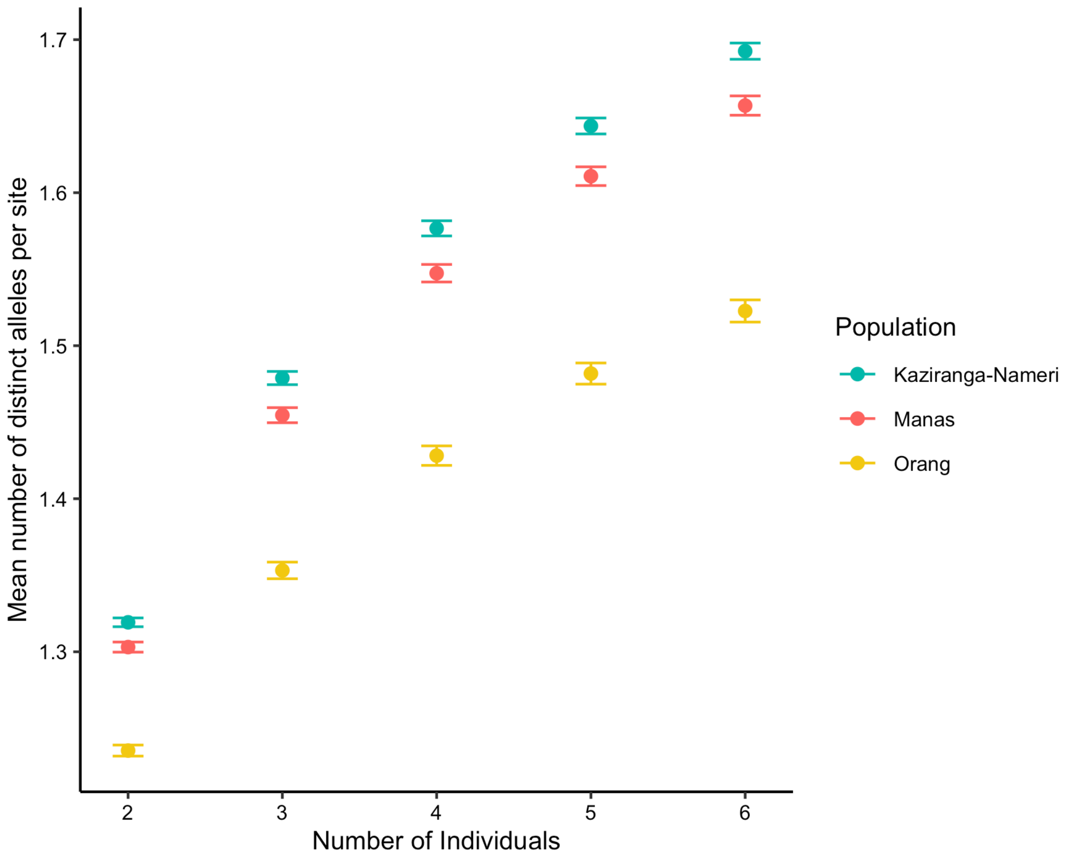


Figure S5: Mean number of distinct alleles per site after correcting for sample size bias. Orang has low genetic variation irrespective of sample size. Pairwise Wilcox test p-value = 0.33 for Orang - Manas, 0.33 for Orang - Kaziranga, and 0.69 for Manas - Kaziranga comparison.

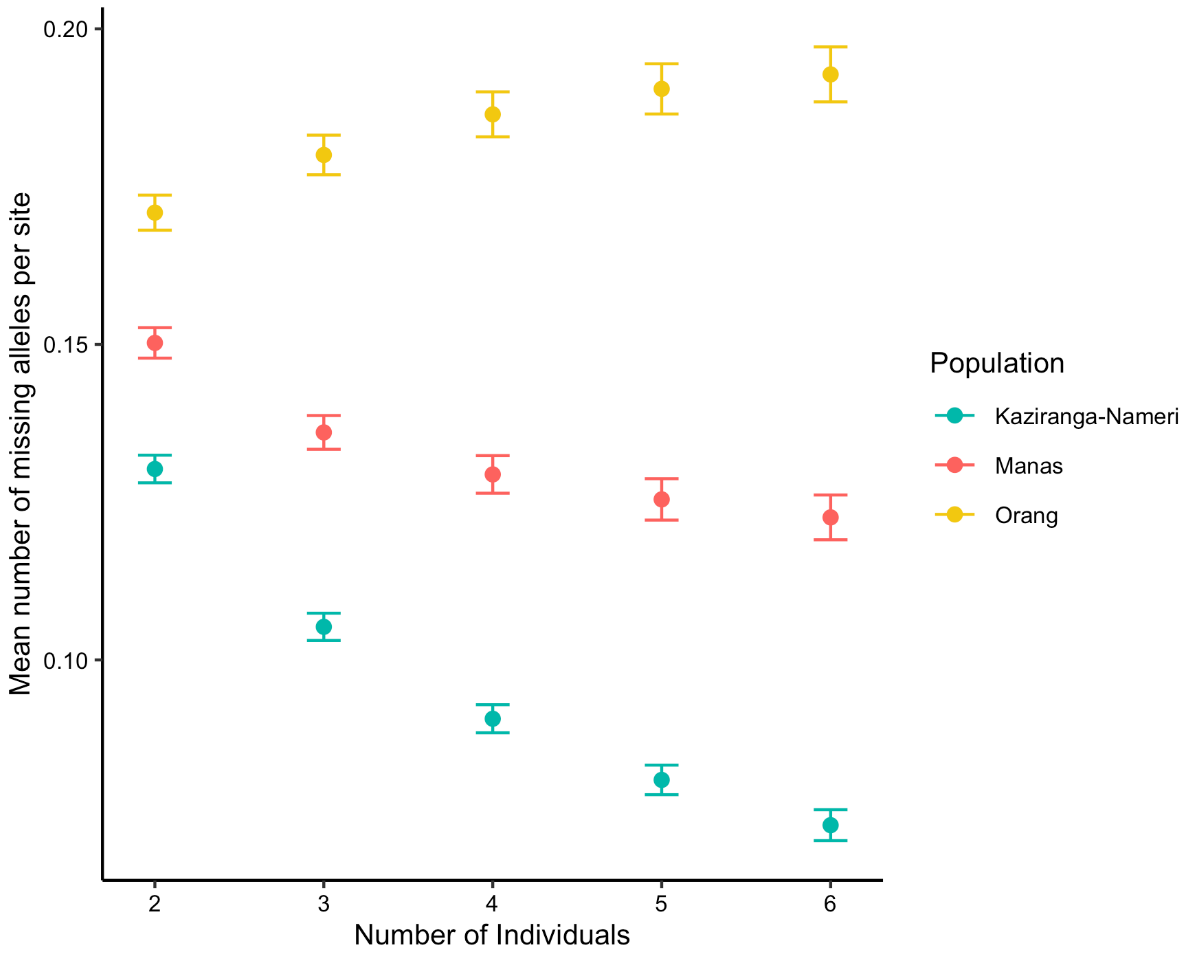


Figure S6: Mean number of alleles private to a pair of PAs which are missing from the third PA after correcting for sampling bias. A large number of alleles are missing from Orang. Assuming the same ancestral population, Orang has lost a lot of genetic variation due to post-bottleneck drift. Also note that Manas has also lost significant variation compared to Kaziranga.

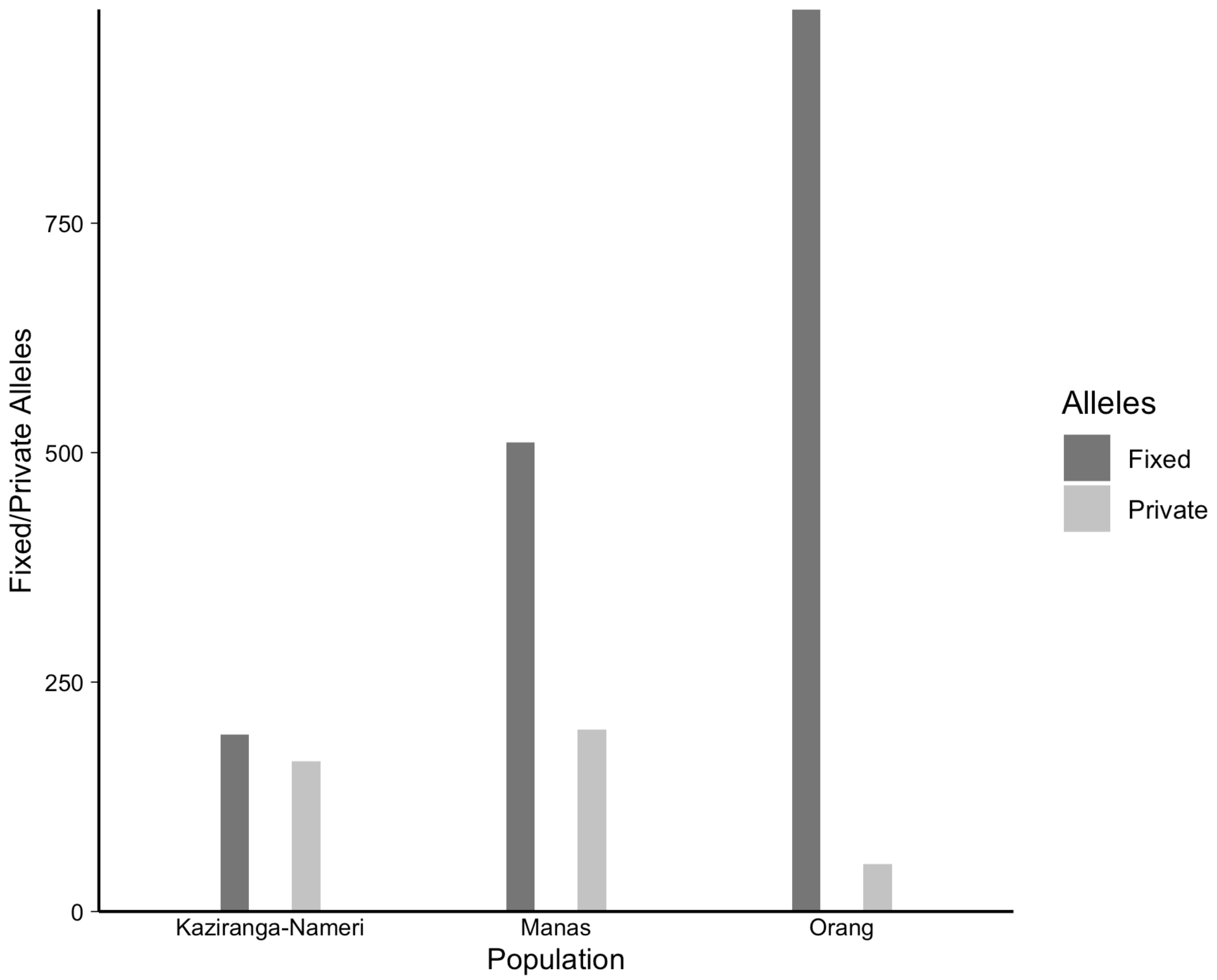


Figure S7: The absolute number fixed and private alleles observed (not corrected for sample size bias). A large number of alleles are fixed in Orang. Orang tigers also have the lowest number of private alleles.

**Supplementary Table 1: Population-wise Downsizing of samples at each step**

| Field Season | Category | Kaziranga | Manas | Orang | Nameri | All pops |
|---|---|---|---|---|---|---|
| **2021** | Total Samples | 133 | 78 | 35 | 21 | 267 |
| | Tigers | 128 | 61 | 35 | 9 | 233 |
| | Tigers% | 96.24 | 78.21 | 100 | 42.86 | 87.27 |
| | Enriched | 37 | 15 | 35 | 9 | 96 |
| | Enriched%(of tigers) | 28.91 | 24.59 | 100 | 100 | 41.20 |
| | Library prepared | 36 | 15 | 21 | 9 | 81 |
| | Library prepared%(of tigers) | 28.13 | 24.59 | 60 | 100 | 34.76 |
| | Sequenced | 16 | 13 | 7 | 8 | 44 |
| | Sequenced%(of tigers) | 12.50 | 21.31 | 20 | 88.89 | 18.88 |
| **2023** | Total Samples | 307 | 119 | 33 | 15 | 474 |
| | Tigers | 282 | 97 | 33 | 9 | 421 |
| | Tigers% | 91.86 | 81.51 | 100 | 60 | 88.82 |
| | Enriched | 170 | 97 | 33 | 9 | 309 |
| | Enriched%(of tigers) | 60.28 | 100 | 100 | 100 | 73.40 |
| | Library prepared | 133 | 73 | 33 | 9 | 248 |
| | Library prepared%(of tigers) | 47.16 | 75.26 | 100 | 100 | 58.91 |
| | Sequenced | 67 | 29 | 29 | 7 | 132 |
| | Sequenced%(of tigers) | 23.76 | 52.58 | 87.88 | 77.78 | 36.58 |
| **Combined** | Total Samples | 440 | 197 | 68 | 36 | 741 |
| | Tigers | 410 | 158 | 68 | 18 | 654 |
| | Tigers% | 93.18 | 80.20 | 100 | 50 | 88.26 |
| | Enriched | 207 | 112 | 68 | 18 | 405 |

| | | | | | | |
|---|---|---|---|---|---|---|
| | Enriched%(of tigers) | 50.49 | 70.89 | 100 | 100 | 61.93 |
| | Library prepared | 169 | 88 | 54 | 18 | 329 |
| | Library prepared%(of tigers) | 41.22 | 55.70 | 79.41 | 100 | 50.31 |
| | Sequenced | 83 | 42 | 36 | 15 | 176 |
| | Sequenced%(of tigers) | 20.24 | 18.35 | 52.94 | 83.33 | 30.28 |

**Supplementary Table 2: Variant sites and samples in the VCF at various filtering steps**

| Filter | Variant sites | Samples |
|---|---|---|
| Raw | 385081559 | 176 |
| Filter 1 | 851617 | 176 |
| Filter 2 | 61293 | 176 |
| Samples with >= 2000 sites; max-miss 0.7; hwe 0.05 | 3226 | 57 |
| Removal of recaptures | 3091 | 44 |

Supplementary Table 3: Estimates of effective population size (Ne) for the four PAs using three different methods implemented in NeEstimator V2 software

| Population | Sample size | Harmonic mean sample size | Non polymorphic sites | Ne | 95% CI (Parametric) | JackKnife on Samples / sites* |
|---|---|---|---|---|---|---|
| Linkage Disequilibrium (LD) Method | | | | | | |
| Kaziranga | 22 | 15.8 | 261 | 30.4 | 30.0 - 30.8 | 10.8 - ∞ |
| Manas | 11 | 9 | 524 | 11.9 | 11.7 - 12.1 | 2.1 - ∞ |
| Nameri | 3 | 1.5 | 1867 | ∞ | ∞ | 0 - ∞ |
| Orang | 8 | 6.4 | 1012 | 1.3 | 1.3 - 1.3 | 0.5 - 15.1 |
| Heterozygote Excess Method | | | | | | |
| Kaziranga | 22 | 18 | 261 | 10.0 | 8.8 - 11.6 | |
| Manas | 11 | 9.5 | 524 | 7.6 | 6.7 - 8.7 | |
| Nameri | 3 | 1.7 | 1867 | 7.6 | 6.2 - 10.0 | |
| Orang | 8 | 6.9 | 1012 | 32.2 | 18.2 - 154.6 | |
| Molecular Coancestry Method | | | | | | |
| Kaziranga | 22 | 18 | 261 | 6.3 | | 5.8 - 6.8 |
| Manas | 11 | 9.5 | 524 | 4.2 | | 4.0 - 4.5 |
| Nameri | 3 | 1.7 | 1867 | 90.2 | | 58.1 - 129.3 |
| Orang | 8 | 6.9 | 1012 | 1.5 | | 1.4 - 1.5 |

Note - Given the low sample size and high number of markers, the LD method is the most accurate and produces estimates close to the true Ne. Both heterozygote excess and molecular ancestry performed poorly because of the inadequate sample size. Notably, the molecular coancestry method, which was suggested to work well for a small inbred population also estimates extremely low Ne for Orang which is very close to the value estimated by the LD method. The LD method fails to estimate Ne for a very low sample size Nameri population, and produces Ne = ∞, implying there is no evidence for variation in the genetic characteristic caused by genetic drift due to a finite number of parents — it can all be explained by sampling error. The given Ne values are for critical frequency 0.05, i.e., any allele with frequency < 0.05 will be removed (this was done to remove singletons).

*JackKnife on samples in the LD method and sites in the molecular coancestry method

Supplementary Table 4: Total and mapped reads for all samples

| Sample | TotalReads | MappedReads | MappingPercentage |
|---|---|---|---|
| KAZ21F008 | 5798755 | 30241 | 0.52 |
| KAZ21F026 | 8571844 | 963282 | 11.24 |
| KAZ21F028 | 5141125 | 5421 | 0.11 |
| KAZ21F033 | 8282473 | 4185957 | 50.54 |
| KAZ21F035 | 7472341 | 15591 | 0.21 |
| KAZ21F036 | 17301751 | 3129651 | 18.09 |
| KAZ21F039 | 7593931 | 9407 | 0.12 |
| KAZ21F004 | 14992251 | 85863 | 0.57 |
| KAZ21F107 | 12217074 | 1879448 | 15.38 |
| KAZ21F125 | 5679588 | 4030 | 0.07 |
| KAZ21F132 | 9846011 | 3512621 | 35.68 |
| KAZ21F013 | 17713750 | 14527816 | 82.01 |
| KAZ21F042 | 14636807 | 45557 | 0.31 |
| KAZ21F057 | 14712255 | 1005893 | 6.84 |
| KAZ21F062 | 15631745 | 147737 | 0.95 |
| KAZ21F076 | 3863892 | 206922 | 5.36 |
| KAZ23F007 | 8900268 | 1520032 | 17.08 |
| KAZ23F011 | 15168792 | 32092 | 0.21 |
| KAZ23F013 | 11096554 | 7319670 | 65.96 |
| KAZ23F019 | 10972759 | 5356909 | 48.82 |
| KAZ23F025 | 18458360 | 43646 | 0.24 |
| KAZ23F028 | 15022661 | 417541 | 2.78 |
| KAZ23F030 | 18847817 | 36699 | 0.19 |
| KAZ23F031 | 8743743 | 29627 | 0.34 |
| KAZ23F032 | 1755860 | 6062 | 0.35 |
| KAZ23F036 | 9820718 | 20808 | 0.21 |
| KAZ23F052 | 9482282 | 61532 | 0.65 |
| KAZ23F057 | 9942426 | 2492568 | 25.07 |
| KAZ23F059 | 10718191 | 11329 | 0.11 |

| | | | |
|---|---|---|---|
| KAZ23F066 | 10657072 | 7900666 | 74.14 |
| KAZ23F078 | 13075175 | 10208295 | 78.07 |
| KAZ23F082 | 7731028 | 200332 | 2.59 |
| KAZ23F096 | 7372503 | 2037 | 0.03 |
| KAZ23F099 | 13270955 | 13495 | 0.1 |
| KAZ23F101 | 22884258 | 6574 | 0.03 |
| KAZ23F103 | 27045858 | 18256 | 0.07 |
| KAZ23F104 | 26329237 | 25829 | 0.1 |
| KAZ23F105 | 22987862 | 26786 | 0.12 |
| KAZ23F106 | 7730798 | 65876 | 0.85 |
| KAZ23F108 | 9953058 | 56624 | 0.57 |
| KAZ23F110 | 12125481 | 2229903 | 18.39 |
| KAZ23F113 | 7461284 | 202660 | 2.72 |
| KAZ23F115 | 7238272 | 123276 | 1.7 |
| KAZ23F116 | 23235260 | 21285102 | 91.61 |
| KAZ23F123 | 15965283 | 173477 | 1.09 |
| KAZ23F127 | 8008052 | 699180 | 8.73 |
| KAZ23F128 | 16072398 | 267560 | 1.66 |
| KAZ23F133 | 14207338 | 19636 | 0.14 |
| KAZ23F135 | 9070168 | 3800 | 0.04 |
| KAZ23F136 | 6264773 | 20045 | 0.32 |
| KAZ23F141 | 15228763 | 68971 | 0.45 |
| KAZ23F145 | 6481200 | 575634 | 8.88 |
| KAZ23F146 | 10809020 | 112278 | 1.04 |
| KAZ23F148 | 16452449 | 1937821 | 11.78 |
| KAZ23F161 | 5957113 | 14709 | 0.25 |
| KAZ23F174 | 20662894 | 74066 | 0.36 |
| KAZ23F187 | 10025254 | 812104 | 8.1 |
| KAZ23F188 | 15986942 | 258358 | 1.62 |
| KAZ23F191 | 7611503 | 273543 | 3.59 |

| | | | |
|---|---|---|---|
| KAZ23F201 | 9642895 | 41487 | 0.43 |
| KAZ23F204 | 13907608 | 20068 | 0.14 |
| KAZ23F209 | 9758915 | 616133 | 6.31 |
| KAZ23F221 | 14339513 | 73107 | 0.51 |
| KAZ23F228 | 14876587 | 8229 | 0.06 |
| KAZ23F230 | 12351765 | 50731 | 0.41 |
| KAZ23F233 | 19379617 | 47631 | 0.25 |
| KAZ23F244 | 13101538 | 26164 | 0.2 |
| KAZ23F245 | 2243912 | 24484 | 1.09 |
| KAZ23F250 | 17320116 | 872504 | 5.04 |
| KAZ23F252 | 2992067 | 4593 | 0.15 |
| KAZ23F263 | 16451812 | 54100 | 0.33 |
| KAZ23F265 | 13165646 | 1830546 | 13.9 |
| KAZ23F266 | 18247067 | 15613 | 0.09 |
| KAZ23F267 | 12743931 | 448177 | 3.52 |
| KAZ23F278 | 7063855 | 8971 | 0.13 |
| KAZ23F289 | 6032854 | 12146 | 0.2 |
| KAZ23F290 | 11635695 | 23291 | 0.2 |
| KAZ23F302 | 9812842 | 1868950 | 19.05 |
| KAZ23F303 | 11318259 | 2384847 | 21.07 |
| KAZ23F304 | 7983189 | 499961 | 6.26 |
| KAZ23F305 | 10625519 | 287545 | 2.71 |
| KAZ23F306 | 14820694 | 34638 | 0.23 |
| KAZ23F307 | 10976491 | 28417 | 0.26 |
| MAN21F02 | 10457430 | 312288 | 2.99 |
| MAN21F21 | 7635420 | 244706 | 3.2 |
| MAN21F30 | 8676925 | 69685 | 0.8 |
| MAN21F38 | 8622272 | 6968290 | 80.82 |
| MAN21F39 | 10228855 | 48037 | 0.47 |
| MAN21F40 | 16536645 | 1996859 | 12.08 |

| | | | |
|---|---|---|---|
| MAN21F41 | 7410768 | 146536 | 1.98 |
| MAN21F44 | 12512404 | 5575884 | 44.56 |
| MAN21F45 | 12855882 | 8305932 | 64.61 |
| MAN21F49 | 11329849 | 1228281 | 10.84 |
| MAN21F55 | 14671766 | 11728508 | 79.94 |
| MAN21F68 | 22498041 | 2440209 | 10.85 |
| MAN21F78 | 17290139 | 12611 | 0.07 |
| MAN23F002 | 7703148 | 14878 | 0.19 |
| MAN23F003 | 8969283 | 3440375 | 38.36 |
| MAN23F006 | 10670906 | 15494 | 0.15 |
| MAN23F009 | 13157021 | 15903 | 0.12 |
| MAN23F010 | 13827801 | 17239 | 0.12 |
| MAN23F012 | 11032213 | 6953623 | 63.03 |
| MAN23F013 | 8161612 | 17558 | 0.22 |
| MAN23F024 | 11790220 | 10074 | 0.09 |
| MAN23F027 | 2630444 | 4424 | 0.17 |
| MAN23F030 | 8847747 | 2137581 | 24.16 |
| MAN23F032 | 10094990 | 56796 | 0.56 |
| MAN23F033 | 9205411 | 5801 | 0.06 |
| MAN23F035 | 9314773 | 22135 | 0.24 |
| MAN23F036 | 13770116 | 9838 | 0.07 |
| MAN23F039 | 6439547 | 16339 | 0.25 |
| MAN23F040 | 10774162 | 44530 | 0.41 |
| MAN23F044 | 5492802 | 282394 | 5.14 |
| MAN23F046 | 14100292 | 1252892 | 8.89 |
| MAN23F047 | 10816857 | 156981 | 1.45 |
| MAN23F050 | 17764418 | 4382 | 0.02 |
| MAN23F051 | 6013266 | 884 | 0.01 |
| MAN23F052 | 5419631 | 63027 | 1.16 |
| MAN23F064 | 8588571 | 14825 | 0.17 |

| | | | |
|---|---|---|---|
| MAN23F068 | 11075129 | 30237 | 0.27 |
| MAN23F070 | 14549684 | 6768 | 0.05 |
| MAN23F076 | 7079705 | 1034597 | 14.61 |
| MAN23F092 | 14038336 | 7690024 | 54.78 |
| MAN23F099 | 11940139 | 12503 | 0.1 |
| MAN23F109 | 4846716 | 5856 | 0.12 |
| NAM21F01 | 9531769 | 107143 | 1.12 |
| NAM21F03 | 9427723 | 11609 | 0.12 |
| NAM21F04 | 8624021 | 595709 | 6.91 |
| NAM21F05 | 13275491 | 91769 | 0.69 |
| NAM21F11 | 680296 | 8148 | 1.2 |
| NAM21F14 | 13021439 | 41923 | 0.32 |
| NAM21F15 | 10315539 | 45857 | 0.44 |
| NAM21F19 | 6763375 | 280783 | 4.15 |
| NAM22F04 | 13748964 | 517984 | 3.77 |
| NAM22F06 | 10794956 | 25944 | 0.24 |
| NAM22F10 | 8973041 | 724683 | 8.08 |
| NAM22F11 | 9442054 | 74072 | 0.78 |
| NAM22F12 | 6272151 | 51963 | 0.83 |
| NAM22F14 | 11595650 | 183252 | 1.58 |
| NAM23F15 | 13095859 | 14107 | 0.11 |
| ORNG21F02 | 12526364 | 550332 | 4.39 |
| ORNG21F06 | 8272755 | 3681733 | 44.5 |
| ORNG21F08 | 8883698 | 102742 | 1.16 |
| ORNG21F12 | 7424940 | 4174448 | 56.22 |
| ORNG21F15 | 7531759 | 9313 | 0.12 |
| ORNG21F20 | 16310485 | 71839 | 0.44 |
| ORNG21F31 | 14677563 | 1420301 | 9.68 |
| ORNG22F01 | 9595972 | 11292 | 0.12 |
| ORNG22F02 | 11923063 | 53311 | 0.45 |

| | | | |
|---|---|---|---|
| ORNG22F03 | 10169174 | 16216 | 0.16 |
| ORNG22F04 | 5946324 | 1526 | 0.03 |
| ORNG22F05 | 12456334 | 8372866 | 67.22 |
| ORNG22F07 | 5226269 | 4634683 | 88.68 |
| ORNG22F08 | 25470124 | 445402 | 1.75 |
| ORNG22F09 | 18535458 | 25284 | 0.14 |
| ORNG22F10 | 9969529 | 6994123 | 70.16 |
| ORNG22F11 | 8974279 | 39937 | 0.45 |
| ORNG22F12 | 9159383 | 100031 | 1.09 |
| ORNG22F13 | 1716672 | 813560 | 47.39 |
| ORNG22F15 | 4521285 | 2167593 | 47.94 |
| ORNG22F16 | 8715022 | 14480 | 0.17 |
| ORNG22F18 | 7997267 | 30349 | 0.38 |
| ORNG22F19 | 2710659 | 370339 | 13.66 |
| ORNG22F21 | 9822831 | 6490007 | 66.07 |
| ORNG22F22 | 6355689 | 5219 | 0.08 |
| ORNG22F23 | 5984846 | 1066 | 0.02 |
| ORNG22F24 | 18202213 | 15071507 | 82.8 |
| ORNG22F25 | 13886305 | 12214605 | 87.96 |
| ORNG22F26 | 16000757 | 142897 | 0.89 |
| ORNG22F27 | 12523099 | 111803 | 0.89 |
| ORNG22F28 | 12026218 | 23836 | 0.2 |
| ORNG22F29 | 9687759 | 257433 | 2.66 |
| ORNG22F30 | 11234638 | 460606 | 4.1 |
| ORNG22F31 | 7996215 | 212427 | 2.66 |
| ORNG22F32 | 4962796 | 2302 | 0.05 |
| ORNG22F33 | 16068371 | 51365 | 0.32 |

Supplementary Table 5: Univariate optimisation

| Surface | Average AIC | Average AICc | Average weight | Average rank | Average R2m | Average LL | Average RMSE | n | Percent top | k |
|---|---|---|---|---|---|---|---|---|---|---|
| NDVI1km | 7079.962708 | 7081.39128 | 0.455822198 | 2.065 | 0.06186586499 | -3535.981354 | 163.5829541 | 504 | 50.4 | 4 |
| Re_LD1km | 7080.320949 | 7081.74952 | 0.3072114157 | 2.065 | 0.178667954 | -3536.160475 | 163.3669443 | 338 | 33.8 | 4 |
| build1km | 7081.308237 | 7082.736808 | 0.185863756 | 2.524 | 0.05440272858 | -3536.654118 | 163.7655857 | 156 | 15.6 | 4 |
| Distance | 7086.571269 | 7086.971269 | 0.03483671522 | 4.16 | 0.04777321979 | -3541.285635 | 165.220592 | 2 | 0.2 | 2 |
| WaterArea1km | 7087.349498 | 7088.77807 | 0.01393939556 | 4.395 | 0.05668535577 | -3539.674749 | 164.9290927 | 0 | 0 | 4 |
| mTPI1km | 7091.163336 | 7092.591907 | 0.002326519502 | 5.791 | 0.04749815758 | -3541.581668 | 165.3166852 | 0 | 0 | 4 |
| Re_LD2km | 7076.538698 | 7077.967269 | 0.5467883166 | 1.464 | 0.1393609421 | -3534.269349 | 163.0740933 | 602 | 60.2 | 4 |
| NDVI2km | 7080.3221 | 7081.750672 | 0.3244891433 | 2.256 | 0.06421778934 | -3536.16105 | 163.7485751 | 341 | 34.1 | 4 |
| build2km | 7080.824708 | 7082.253279 | 0.1096943653 | 2.664 | 0.05555872583 | -3536.412354 | 163.6582737 | 57 | 5.7 | 4 |
| Distance | 7088.127505 | 7088.527505 | 0.01183418737 | 4.274 | 0.04733150719 | -3542.063753 | 165.4392631 | 0 | 0 | 2 |
| WaterArea2km | 7088.692698 | 7090.121269 | 0.006424990673 | 4.529 | 0.05819707959 | -3540.346349 | 165.130765 | 0 | 0 | 4 |
| mTPI2km | 7092.874068 | 7094.302639 | 0.0007689967772 | 5.813 | 0.04712181588 | -3542.437034 | 165.5607343 | 0 | 0 | 4 |
| build5km | 7073.658842 | 7075.087413 | 0.5201803537 | 1.782 | 0.06125678363 | -3532.829421 | 162.5532498 | 557 | 55.7 | 4 |
| Re_LD5km | 7076.003724 | 7077.432296 | 0.2789387069 | 1.936 | 0.06649939388 | -3534.001862 | 163.1466539 | 287 | 28.7 | 4 |
| NDVI5km | 7081.629077 | 7083.057648 | 0.05233231726 | 3.497 | 0.05909687536 | -3536.814538 | 163.7564327 | 16 | 1.6 | 4 |
| WaterArea5km | 7082.82857 | 7084.257141 | 0.1315848922 | 3.639 | 0.05277191189 | -3537.414285 | 164.1730109 | 139 | 13.9 | 4 |
| Distance | 7087.419599 | 7087.819599 | 0.01593330474 | 4.427 | 0.04959366077 | -3541.7098 | 165.310585 | 1 | 0.1 | 2 |
| mTPI5km | 7092.346721 | 7093.775293 | 0.001030425177 | 5.719 | 0.04935046026 | -3542.173361 | 165.460073 | 0 | 0 | 4 |
| build10km | 7052.724064 | 7054.152635 | 0.9658713266 | 1.02 | 0.08435439761 | -3522.362032 | 159.4749579 | 988 | 98.8 | 4 |
| NDVI10km | 7067.069314 | 7068.497886 | 0.01070180399 | 2.499 | 0.06829367833 | -3529.534657 | 161.5164984 | 2 | 0.2 | 4 |
| WaterArea10km | 7070.629416 | 7072.057987 | 0.007296234122 | 3.328 | 0.05135177532 | -3531.314708 | 162.1907579 | 2 | 0.2 | 4 |
| Distance | 7079.62463 | 7080.02463 | 0.008406800501 | 3.745 | 0.05144414946 | -3537.812315 | 164.0327317 | 1 | 0.1 | 2 |
| mTPI10km | 7081.952137 | 7083.380709 | 0.007043550718 | 4.777 | 0.05572080831 | -3536.976069 | 163.723987 | 7 | 0.7 | 4 |
| Re_LD10km | 7083.62463 | 7085.053202 | 0.0006802841146 | 5.631 | 0.05144414946 | -3537.812315 | 164.0327317 | 0 | 0 | 4 |
| build20km | 7049.740321 | 7051.168893 | 0.9739414208 | 1.008 | 0.08342901081 | -3520.870161 | 158.8891665 | 992 | 99.2 | 4 |
| Re_LD20km | 7065.881281 | 7067.309852 | 0.01511812647 | 2.408 | 0.0518295539 | -3528.94064 | 161.4214016 | 6 | 0.6 | 4 |
| NDVI20km | 7069.530794 | 7070.959366 | 0.002602141995 | 3.383 | 0.06734779587 | -3530.765397 | 161.9162741 | 0 | 0 | 4 |
| WaterArea20km | 7069.821901 | 7071.250473 | 0.006672778974 | 3.518 | 0.05124400894 | -3530.910951 | 162.008316 | 2 | 0.2 | 4 |
| Distance | 7085.425241 | 7085.825241 | 0.001077495895 | 4.816 | 0.04820976734 | -3540.71262 | 165.0213607 | 0 | 0 | 2 |
| mTPI20km | 7088.531021 | 7089.959592 | 0.0005880359038 | 5.867 | 0.04934005234 | -3540.26551 | 164.8547167 | 0 | 0 | 4 |
